# FAD mutations suppress brain angiogenesis and neuroprotection by reducing γ-secretase processing and dimerization of VEGFR2

**DOI:** 10.64898/2026.05.12.724648

**Authors:** Rukmani Pandey, Amira Zarrouk, Poulomi Dey, Elizabeth Levendosky, Gilles Carpentier, Patrick Hof, Anastasios Georgakopoulos, Nikolaos K. Robakis

## Abstract

Efficient cerebrovasculature is vital to neuronal health and cognition and evidence shows that most dementia patients have cerebrovascular abnormalities. Brain vasculature is regulated by Vascular Endothelial Growth Factors (VEGFs) binding VEGF receptor2 (VEGFR2) and stimulating angiogenesis and neuroprotection. Here we show that an ADAM17 cleavage of extracellular VEGFR2 produces the membrane-bound γ-secretase substrate VEGFR2/CTF1 (called VCTF1), comprising the transmembrane and intracellular domains of VEGFR2. VCTF1 binds full-length VEGFR2 monomers suppressing its dimerization a function that is required for VEGFR2 activation and downstream angiogenesis and neuroprotection. Presenilin1 (PS1) Familial Alzheimer’s disease (FAD) mutants exert dominant negative effects on the γ-secretase processing of VCTF1 increasing its concentration and abolishing VEGF-A-induced VEGFR2 dimerization/activation and downstream VEGFR2 signaling, endothelial cell functions and angiogenesis. γ-Secretase inhibitors or PS1 reduction have similar effects on VCTF1 accumulation and VEGFR2 dimerization/activation and downstream signaling and functions as PS1 FAD mutants. Moreover, PS1 FAD mutants increase vulnerability of brain neurons to ischemic stress and abolish VEGF-A-induced neuroprotection and cognition. Together, these data show that VCTF1 suppresses VEGFR2 dimerization and downstream signaling and functions of the brain’s VEGF-A-/VEGFR2 angiogenic and neuroprotection systems. Importantly, we detected molecular markers of decreased VEGFR2 dimerization and angiogenic dysfunction in human brain tissue from PS1 FAD mutant genotypes. Our data reveal a pathway through which FAD mutants may promote dementia by increasing accumulation of VCTF1 and decreasing angiogenesis, neuroprotection, and cognition, suggesting that PS1 FAD patients may benefit from therapeutic methods that decrease brain VCTF1.

## Introduction

The cerebrovascular system is essential for neuronal survival and function and extensive research shows that cerebrovascular abnormalities are frequently present in patients with Alzheimer’s disease (AD) and other forms of dementia [1–3]. Notably, AD is associated with a higher prevalence of cerebrovascular abnormalities than other neurodegenerative disorders [4], and cerebrovascular dysfunction is a significant risk factor for AD [4,5], with estimates suggesting that approximately 75% of AD patients also have brain vascular abnormalities [1,6]. AD neuropathology, including amyloid plaques and neurofibrillary tangles (NFTs), commonly coexists with brain vascular abnormalities [7, 8] in Sporadic (SAD) and Familial AD (FAD) [9–13], and brain microvessel (MV) pathology correlates with cognitive decline in individuals at risk of dementia [7,14]. Vascular dysfunction is an early manifestation of many dementias including AD [15], promoting neurodegeneration via impaired cerebral perfusion [16], metabolic dysregulation [17], and inflammation [15,18]. The brain responds to damaged vasculature by stimulating angiogenesis and neovascularization, promoting tissue repair [19]. For example, ischemia, a vascular pathology commonly observed in SAD [20,21] and FAD brains [13], induces neovascularization by stimulating expression of vascular endothelial growth factors (VEGFs), particularly VEGF-A that binds with high affinity VEGF receptor2 (VEGFR2), the principal endothelial cell (EC) signaling receptor and a key regulator of cerebral angiogenesis and vascular integrity [22,23]. This ligand-receptor interaction stimulates VEGFR2 dimerization [22–24], increasing its autophosphorylation/activation and triggering downstream angiogenic signaling, EC functions, angiogenesis and neuroprotection [22,23]. Impaired reparative angiogenesis can lead to reduced brain blood flow and decreased delivery of oxygen and nutrients, increasing the brain’s vulnerability to injury and neuronal death [5]. Additional studies indicate that vascular risk factors and alterations in cortical blood flow emerge years before the onset of cognitive decline [5,9,25,26]. Notably, elevated VEGFs found in AD brains with vascular pathology, may suggest unsuccessful attempts to activate VEGFR2-mediated neovascularization and neuronal survival [27,28].

VEGFR2 is a type I cell surface transmembrane (TM) receptor tyrosine kinase (RTK) that undergoes extracellular cleavage by metalloproteinase (MP) ADAM17, a process stimulated by VEGF-A [29]. However, it remains unclear whether VEGFR2 is also processed by γ-secretase, a multi-subunit TM aspartyl protease complex known to cleave type I transmembrane proteins with implications for cell signaling and function [30,31]. γ-Secretase contains either Presenilin 1 (PS1) or Presenilin 2 (PS2) as its catalytic subunit, with PS1 being the predominant proteolytic component [30,32]. Although PS1 mutations are the most common cause of FAD (www.alz.org), the mechanisms promoting neurodegeneration in FAD brains remain incompletely understood. Here we show that decreased γ-secretase cleavage of the VEGFR2-derived peptide VCTF1 suppresses VEGF-A-stimulated VEGFR2 dimerization/activation and downstream angiogenesis and neuroprotection. These findings reveal a pathway through which γ-secretase processing of VCTF1 controls brain angiogenesis and a mechanism via which PS1 FAD mutants suppress brain angiogenesis and neuroprotection. Our data suggest that enhancing metabolism of peptide VCTF1 may have therapeutic benefits for FAD patients.

## Results

### 1. PS1 FAD mutants and RO γ-secretase inhibitor (GSI) impair VEGF-A-induced brain angiogenic signaling, EC functions and angiogenesis

Angiogenesis plays critical roles in tissue development and adult wound healing and VEGFR2 is the primary mediator of brain EC proliferation and vascularization [23,25]. VEGFR2-mediated angiogenesis is induced by VEGF-A which plays a primary role in the stimulation of brain angiogenesis [22,23]. To investigate effects of PS1 FAD mutations on brain angiogenesis, we used wild-type (WT) mice and two knock-in (KI) transgenic mouse lines expressing either FAD mutant PS1M146V or FAD mutant PS1I213T [33–36], (see also Methods). In the heterozygous (HTRZ) state, KI mice mirror the HTRZ genetic condition of PS1 FAD patients. To stimulate brain angiogenesis, WT and HTRZ PS1 FAD mutant-expressing mice were subjected to carotid artery infusion of VEGF-A [37] and microvascular remodeling was measured by following microvascular length density [38]. As a second indicator of angiogenesis we followed formation of the VEGFR2/endoglin complex, part of the tripartite VEGFR2-endoglin-neuropilin1 complex that promotes VEGF-A-induced sprouting angiogenesis [39]. Fig. 1A (upper two panels) shows that in contrast to WT mouse brains that increase microvessel density in response to VEGF-A treatment, mice expressing FAD mutant PS1M146V show no stimulation of brain microvessel density in response to VEGF-A, indicating reduced microvascular structural integrity and remodeling in PS1 FAD mice following ischemia, consistent with an impaired angiogenic response. To further assess reproducibility of data obtained with PS1M146V-expressing mice, we tested a second HTRZ KI mouse model carrying FAD mutant PS1I213T and obtained data similar to those obtained with mutant PS1M146V (Fig. 1A, lower panel). Moreover, as shown in Fig. 1B, VEGF-A-induced formation of the VEGFR2/endoglin complex is significantly reduced in mouse brains expressing either one of the above PS1 FAD mutants compared to WT controls further supporting the hypothesis that PS1 FAD mutations impair the VEGF-A-induced brain angiogenesis.

**Figure 1.**
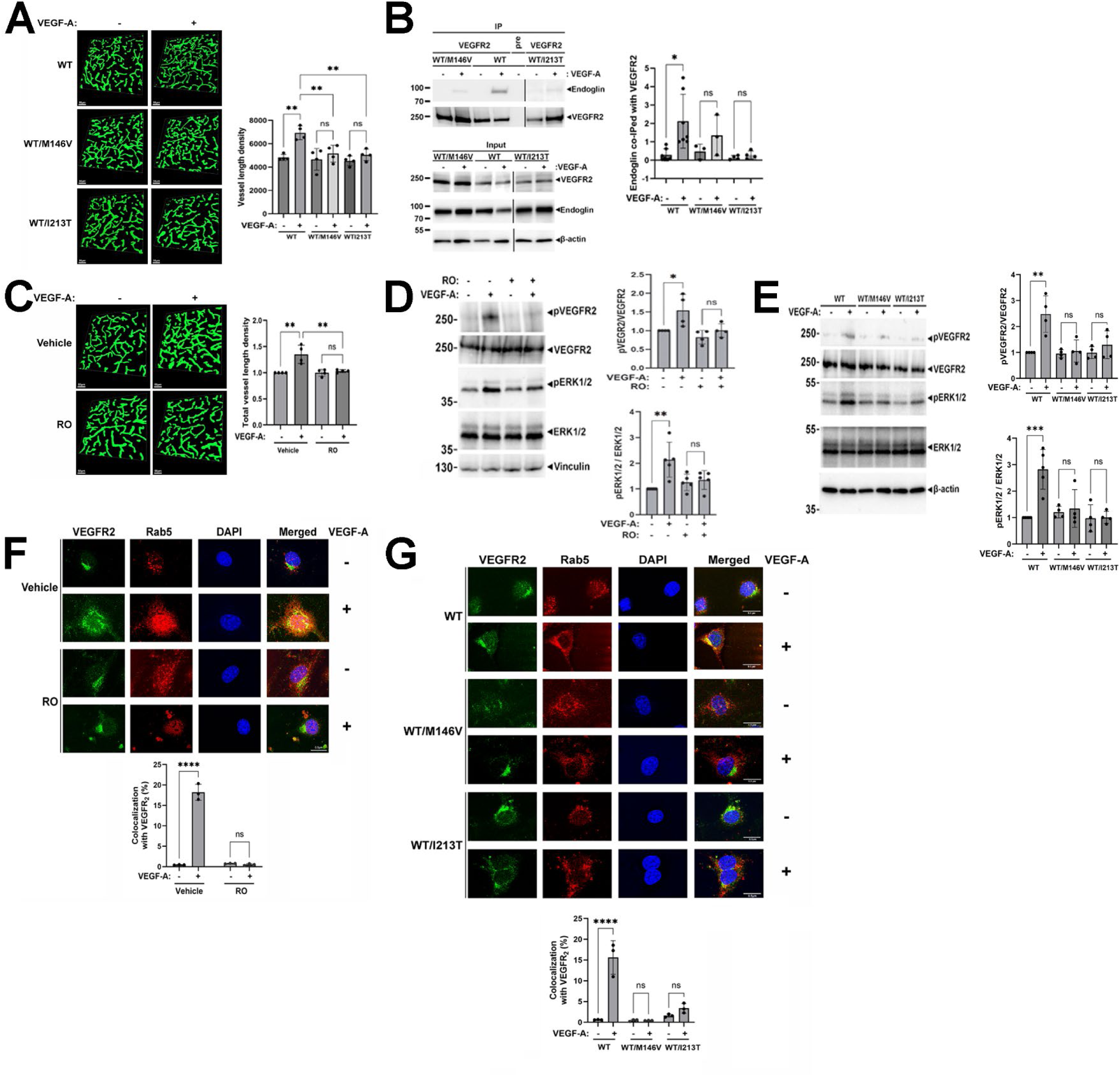
PS1 FAD mutants and γ-secretase inhibitors impair the VEGF-A-stimulated angiogenic signaling, functions and angiogenesis. (A), Wild-type (WT) and KI mice HTRZ for either PS1 FAD mutant M146V (WT/M146V) or I213T (WT/I213T) were infused with vehicle (0.2% BSA in PBS) or VEGF-A (a total of 3.5μg in 100μl of vehicle) through the carotid artery for 15 days using a mini osmotic pump as in Methods. Left: Brain coronal sections (40μm thick) were prepared and immunostained with anti-Col IV antibodies to visualize brain vessels. Enhanced visualization surfaces were generated using Imaris software from representative confocal images of ipsilateral hemispheres. Scale bar: 50μm. Right: Graph shows total vessel length density in WT and PS1 FAD brains quantified using Imaris 9.9 software as in Methods. (B), WT and HTRZ for PS1 FAD mutants M146V or I213T mice were injected through the carotid artery for 20 minutes with either vehicle or 100ng of VEGF-A in vehicle prepared as in 1A using a catheter as described in Methods. Brain microvessels (MV) were isolated as in Methods, lysed in Triton X-100 buffer, and subjected to immunoprecipitation (IP) with anti-VEGFR2 antibody or control IgG. Left: IPs were analyzed on Western blots (WBs) using anti-endoglin or anti-VEGFR2 antibodies (upper panel). Input samples are shown in lower panel. β-actin: loading control. Right: Graph shows quantification of endoglin co-IPed with VEGFR2, normalized to IPed VEGFR2. (C), WT mice were infused for 15 days through the carotid artery with vehicle or VEGF-A in vehicle as in 1A using a mini osmotic pump (as in 1A). For RO injection, mice were treated with vehicle (2% DMSO, 30% PEG 300, 5% Tween-80 in ddH2O) or RO in vehicle (5mg/kg body weight) via five injections in tail vein one injection every three days, with first injection administered 1 hour before osmotic pump implantation. Brain coronal sections (40μm) were prepared and immunostained with anti-Col IV antibodies as in 1A. Left: Representative confocal images of ipsilateral hemispheres are shown prepared as in 1A. Scale bar: 50μm. Right: Graph shows total vessel length density quantified using Imaris software as in 1A. (D), WT adult mice were treated with either 50μl vehicle as in 1C or 1mg/kg RO in vehicle via carotid artery as in Methods. 15-16 hrs later, 50 μl vehicle prepared as in 1A or 100ng VEGF-A in vehicle was administered via carotid artery for 10-20 minutes using a catheter as in 1B. Brain MVs were isolated and extracted as in 1B. Left: p-VEGFR2 (Tyr1054/Tyr1059), VEGFR2, p-ERK1/2 and ERK1/2 are detected on WBs of extracts with specific antibodies in MV extracts. Vinculin: loading control. Right: Graphs show fold change of phosphorylated to total protein ratio. (E), WT mice and mice HTRZ for PS1 FAD mutant M146V (WT/M146V) or I213T (WT/I213T) were treated with vehicle or VEGF-A via carotid artery for 10-20 minutes using a catheter as in 1D. Brain MVs were isolated and extracted as in 1B. Left: p-VEGFR2 (Tyr1054/Tyr1059), VEGFR2, p-ERK1/2 and ERK1/2 are detected on WBs of extracts with specific antibodies in MV extracts. β-actin: loading control. Right: Graphs show fold change of phosphorylated to total protein ratio. (F), WT pCECs were prepared and treated as in Methods with vehicle (DMSO) or RO (200nM in DMSO) and then stimulated with either vehicle (PBS) or VEGF-A (20ng in PBS) for 15min. Upper: Cells were co-immunostained with either anti-VEGFR2 antibodies (green) or early endosome marker Rab5 (red) and cell nuclei were stained with Hoechst (blue) as in Methods. Yellow fluorescence in merged images indicates co-localization of VEGFR2 with Rab5. Scale bar 0.5μm. Lower: Graph shows percent of VEGFR2 co-localized with Rab5 in RO-treated WT cells compared to vehicle-treated cells measured with Imaris software. (G), pCECs from either WT or mice HTRZ for PS1 FAD mutant M146V (WT/M146V) or I213T (WT/I213T), were stimulated with vehicle or VEGF-A in vehicle as in 1F. Upper: Cells were co-stained with anti-VEGFR2 antibodies and early endosome marker Rab5 as in 1F. Cell nuclei were stained with Hoechst (blue) as in 1F. Yellow fluorescence in merged images indicates co-localization of VEGFR2 with Rab5. Scale bar 0.5μm. Lower: Graph shows percent of VEGFR2 co-localized with Rab5 in PS1 FAD WT/M146V or WT/I213T HTRZ mice compared to WT measured with Imaris software. For Figs A-G, data are shown as Mean ± S.E. from at least three independent experiments or as indicated in the dot plots. Statistical analysis was performed using two-way ANOVA followed by Tukey post-hoc test. ns = not significant, *p<0.05, **p<0.01, ***p<0.001.

PS1 FAD mutants have been shown to reduce the γ-secretase cleavage activity at the epsilon (ε) site of substrates [40–43] raising the possibility that such mutants suppress VEGF-A-induced brain angiogenesis by decreasing the γ-secretase activity of PS1. As GSIs may have pleiotropic effects on angiogenesis, stimulating or decreasing it depending on model and experimental conditions [44,45], we used RO4929097 (RO) [33,46], a potent and selective GSI that has consistently shown inhibitory effects on angiogenesis. Due to its high selectivity and ability to strongly deactivate Notch receptors, GSI RO was used in anti-cancer clinical trials [46–48]. Fig. 1C shows that similar to effects of PS1 FAD mutants on brain vascular density, treatment of WT mice with RO also reduces VEGF-A-stimulated brain vascular density compared to mock-treated control mice. Under our conditions, neither PS1 FAD mutants [33] nor the RO inhibitor affected the viability of ECs (data not shown and ref. 46). These findings support the suggestion that γ-secretase activity is required for VEGF-A-induced brain angiogenesis. Moreover, similar to RO, PS1 FAD mutants suppress cleavage of γ-secretase substrates [40–43] raising the possibility that FAD mutants reduce brain angiogenesis by decreasing processing of γ-secretase substrates.

VEGF-A-induced angiogenesis depends on VEGF-A binding to EC surface VEGFR2 stimulating its dimerization and autophosphorylation, a process that in turn activates downstream VEGFR2 signaling steps crucial to angiogenesis such as internalization and trafficking of VEGFR2 to endosomes [23] and activation of survival kinases including ERK1/2 that is essential to EC migration and proliferation [23]. To test the importance of γ-secretase activity on the VEGF-A-induced autophosphorylation/ activation of brain VEGFR2 (pVEGFR2) and phosphoERK1/2 kinase (pERK1/2), we treated WT mice with GSI RO followed by stimulation with VEGF-A (see Methods). Mock-treated mice served as controls and brain MVs were prepared from all samples and extract analyzed on western blots (WBs) as in Methods. Fig. 1D (two left lanes) shows that in the absence of RO treatment, injection of VEGF-A via the mouse carotid stimulates phosphorylation of both pVEGFR2 at residues 1054/1059 [49] and pERK1/2 in brain MVs. In contrast, RO treatment suppresses the VEGF-A-stimulated production of pVEGFR2 and pERK1/2 in brain MVs (Fig. 1D, two right lanes). Furthermore, compared to WT (Fig. 1E, two left lanes), brain MVs from mice HTRZ for PS1 mutant M146V or I213T treated with VEGF-A contain reduced levels of both pVEGFR2 (residues 1054/1059) and pERK1/2 (Fig. 1E, four right lanes). Together, these data show that RO and PS1 FAD mutants have similar negative effects on the VEGF-A-induced phosphorylation /activation of pVEGFR2 and pERK1/2. Using mouse brain primary Cortical EC Cultures (pCECs), we obtained similar data for the *<u>in vitro</u>* effects of RO inhibitor and PS1M146V mutant on the VEGF-A-stimulated pVEGFR2 and pERK1/2 (Suppl. Figs. 1A, B), providing further support for the hypothesis that both, RO and PS1 FAD mutants reduce the VEGF-A-stimulated angiogenic signaling to VEGFR2 and ERK1/2.

VEGF-A-stimulated internalization and trafficking of cell surface VEGFR2 to early and late endosomes and lysosomes are key steps for downstream angiogenic signaling [23,50]. We found that treatment of WT pCECs with RO (Fig. 1F) or pCECs from mice HTRZ for PS1 FAD mutants (Fig. 1G), reduce VEGF-A-induced trafficking of VEGFR2 to early (Figs. 1F,G) and late (Suppl. Fig. 1C,D) endosomes marked by Rab5 or Rab7 respectively. In addition, compared to controls, pCECs treated with RO inhibitor or expressing FAD mutants reduce VEGFR2 trafficking to LAMP-2-expressing lysosomes (Suppl. Figs.1E,F). Together, these findings show that both RO and PS1 FAD mutants impair VEGF-A-stimulated angiogenic signaling steps such as phosphorylation/activation of VEGFR2 and ERK1/2 kinase and intracellular trafficking of VEGFR2. Moreover, our data suggest that RO and FAD mutants impair angiogenesis by targeting a step upstream of the pVEGFR2 activation of the angiogenic cascade [22,23].

VEGF-A-induced EC functions including sprouting, capillary tube formation, and cell migration are crucial to the initiation of angiogenesis and neovascularization as their detection indicates that ECs are primed to initiate the process of angiogenesis [33,51]. Figures 2A–C show that VEGF-A-induced sprouting, migration, and tube formation steps are significantly attenuated in pCECs expressing PS1 FAD mutant M146V or I213T, compared to those of WT pCECs. In addition, GSI RO impairs the VEGF-A-stimulated sprouting, migration, or tube formation of WT pCECs (Figs. 2D-F). These findings show that both GSI RO and PS1 FAD mutants suppress VEGF-A-stimulated brain angiogenesis, angiogenic signaling, and EC functions. Together, our data indicate that γ-secretase activity is required for brain angiogenesis, and futher suggest that suppression of VEGF-A-stimulated angiogenesis by PS1 FAD mutations may be due to the reduced γ-secretase activity of PS1 FAD mutants.

**Figure 2.**
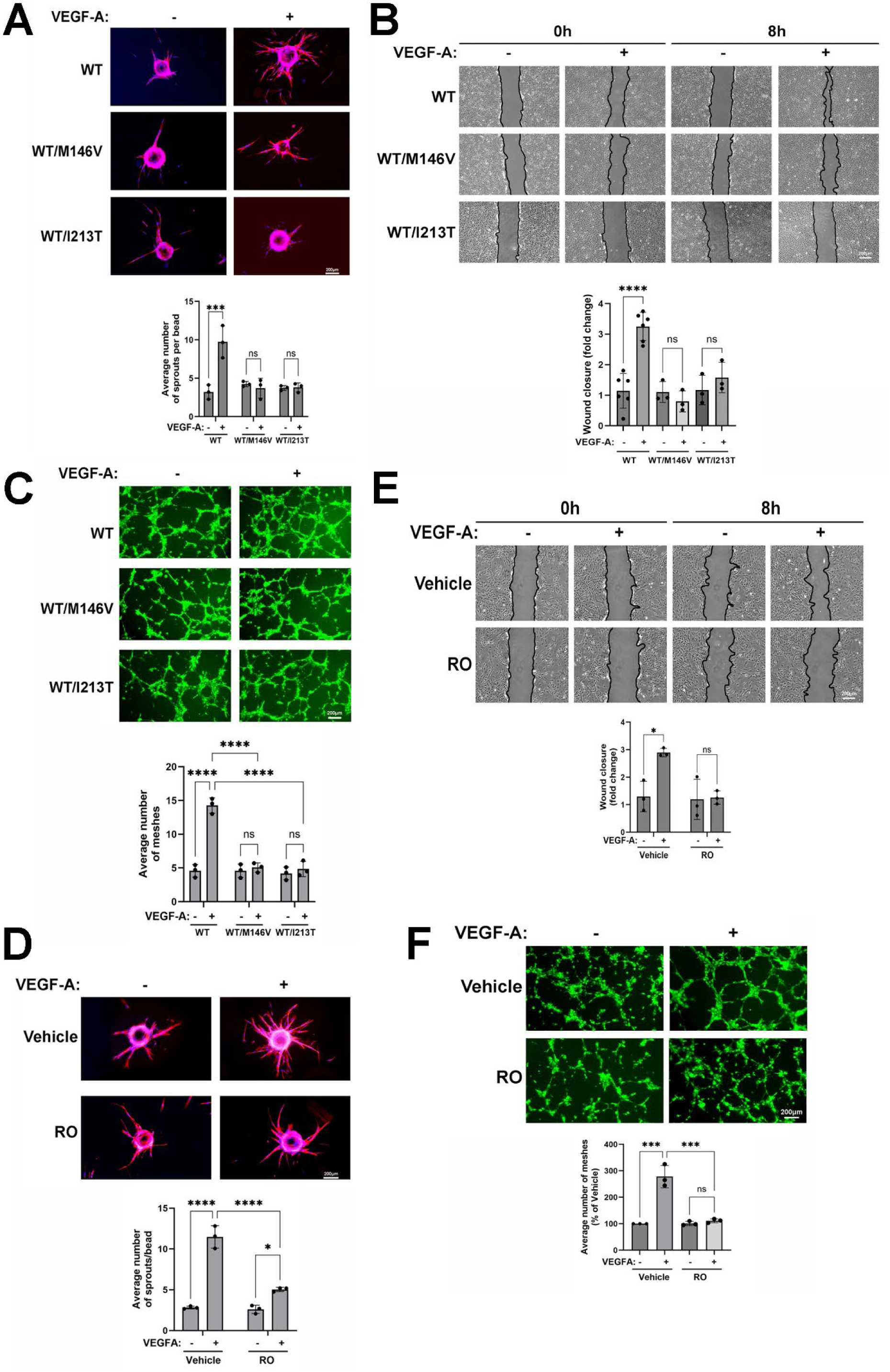
PS1 FAD mutants and γ-secretase inhibitor decrease VEGF-A-induced EC functions. pCECs from WT, WT/M146V- or WT/I213T-expressing mice were prepared as in Methods. **(A),** Cells grown on microcarrier beads were treated with either vehicle (PBS) or VEGF-A (10ng/ml in vehicle) for 72 hours. **Upper panels:** representative fluorescent photomicrograph showing Hoechst-stained nuclei (blue) and phalloidin-labelled actin cytoskeleton (red). **Lower:** Graph shows quantification of sprouting as average number of sprouts per bead. **(B),** pCECs were seeded in 35-mm dishes, scratched to create wounds on a confluent monolayer and then treated with either vehicle or VEGF-A as in 2A. Migration was assessed as in Methods. **Upper panels:** Representative phase contrast photomicrographs showing wound areas outlined in black. **Lower:** Graph shows quantification of cell migration represented as fold change in wound (scratch) closure relative to WT. **(C),** pCECs were seeded as in Methods and treated as in 2A for 3 hours. Tubes were quantified as in Methods. **Upper panels:** representative photomicrographs (Calcein-AM; green) showing tube-like (loop/mesh) structures. **Lower:** Graph shows quantification of tube formation as average number of loops/meshes per field. **(D-F),** WT pCECs were treated with either vehicle or VEGF-A as in 2A in the presence of either vehicle or RO as in 1F. Graphs show quantification of sprouting **(D)** migration **(E)** and tube formation **(F)**. **A-F:** statistical significance was determined using two-way ANOVA followed by Tukey post-hoc analysis. ns = not significant, *p<0.05, **p<0.01, ***p<0.001.

### 2. PS1 FAD mutants impair the γ-secretase processing of brain VEGFR2

VEGFR2 is a type I TM cell surface RTK mediating VEGF-A-induced sprouting angiogenesis. Mature VEGFR2 protein contains about 1350 amino acids having an SDS-PAGE mobility of about 250 kDa due to a large number of post-translational modifications including glycosylation, phosphorylation and ubiquitination [23,49,52]. In contrast, the apparent MW of VEGFR2 transfected into HEK293T cells is about 180kDa due to reduced posttranslational modifications [53]. VEGFR2 has a soluble extracellular domain of about 760 amino acids, a TM sequence of 24 residues and a cytosolic domain of about 560 residues [49]. Membrane protease ADAM17 cleaves extracellular domains of type I TM proteins near the surface of the plasma membrane [54,55] and VEGF-A stimulates the ADAM-17 cleavage and shedding of the extracellular soluble VEGFR2 [29]. This cleavage also produces a remaining membrane-bound C-terminal fragment1 (CTF1) containing the TM and intracellular domains of VEGFR2, called here VCTF1. A large number of CTF1 fragments of cell surface proteins are processed by γ-secretase [30,31,56], but the fate of the membrane-bound VCTF1 remaining after the ADAM17-mediated cleavage of VEGFR2 is still unknown. VEGF-A-treatment of membranes prepared from cells expressing VEGFR2 tagged at its C-terminus with an eight residue c-Myc tag (VEGFR2-Myc, see Methods), revealed a c-Myc-reactive peptide of SDS-PAGE MW about 72kDa (marked VCTF1-Myc in Fig 3A, lanes 2 and 4) suggesting this peptide is the remaining membrane-bound fragment of VEGFR2 following its cleavage by ADAM17. Using an *<u>in vitro</u>* γ-secretase membrane assay [40], we found that incubation of GSI RO with membranes from cells expressing VEGFR2-Myc accumulates the 72kDa VCTF1-Myc peptide suggesting VCTF1 is a γ-secretase substrate (Fig. 3A, lanes 2 and 3). As expected, the VCTF1-Myc peptide accumulating in response to RO also reacts with antibodies against cytoplasmic VEGFR2 (Fig. 3B, left) and its levels increase by VEGF-A while decreasing by ADAM17 inhibitors (Fig. 3B, right). Treatment of our membrane preparations with VEGF-A also produced an additional c-Myc-reactive peptide of about 65kDa (marked VCTF2-Myc, Fig. 3A, lanes 2 and 4) indicating it contains cytoplasmic sequence of VEGFR2. VCTF2-Myc is not detected in the presence of RO inhibitor (Fig. 3A lanes 4 and 5) strongly suggesting VCTF2 is the expected CTF2 product of the γ-secretase-Regulated Intramembrane Proteolysis (RIP) of peptide VCTF1 [30,31,56]. Together, these findings show that VCTF1 is a membrane-bound γ-secretase substrate derived through the ADAM17 cleavage of VEGFR2. VCTF1 comprises the TM and cytoplasmic domains of VEGFR2, probably along with approximately 15 residues of extracellular VEGFR2 extending from the outer plasma membrane to the consensus sequence of the ADAM17 cleavage of VEGFR2 [55].

**Figure 3.**
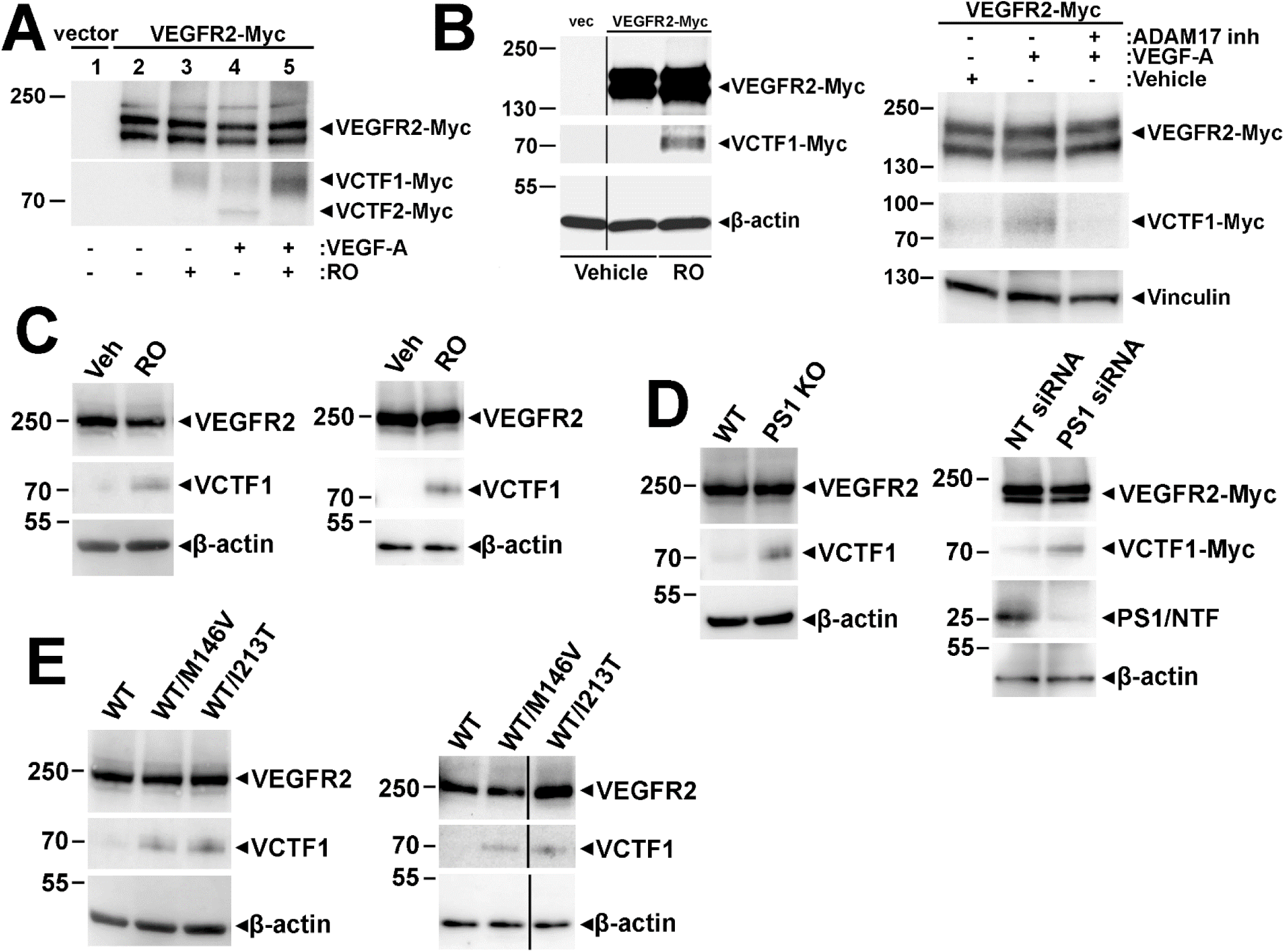
VEGFR2 is processed by PS1/γ-secretase and PS1 FAD mutants decrease processing. **(A),** HEK293T cells transfected with either pCMV3 vector or VEGFR2-Myc-expressing vector were treated with either vehicle or RO overnight as in 1F. Membrane fractions were prepared as in Methods and incubated with vehicle (-) or VEGF-A as in 1F for 30 minutes in the presence of lactacystin. VEGFR2-Myc, VCTF1-Myc, and VCTF2-Myc, were then detected on WB using anti-Myc antibodies. Representative blot shows Myc-labelled fragments as indicated in Figure. **(B), Left:** HEK293T cells were transfected with VEGFR2-Myc and treated with vehicle or RO as in 3A and extracted in SDS buffer as in methods. VEGFR2-Myc and VCTF1-Myc were detected in cell extracts on WBs with antibodies recognizing the cytoplasmic sequence of VEGFR2 (ab39256). β-actin: loading control. **Right:** HEK293T cells were transfected as in 3A. Cells were pretreated with 200nm ADAM17 inhibitor D1 (A12: ADAM17 inh) for 1 hour and then stimulated with vehicle or VEGF-A as in 3A for 1h. Cells were extracted as in 3B Left. VEGFR2-Myc and its proteolytic product VCTF1-Myc are detected with anti-Myc antibody on WB as in 3A. Vinculin: loading control. **(C), Left:** WT pCECs were treated with either vehicle (DMSO; Veh) or RO as in 1F for 15-16 h and extracted in SDS buffer. VEGFR2 and VCTF1 were detected in cell extracts on WB with anti-VEGFR2 antibody as in 3B Left. β-actin: loading control. **Right:** Brain MVs were isolated from adult WT mice as in 1B following 15 hours of treatment with either vehicle or RO as in 1D and extracted in SDS buffer as in Methods. VEGFR2 and VCTF1 were detected in MV extracts on WB with anti-VEGFR2 antibody as in 3B Left. β-actin: loading control. **(D), Left:** pCECs isolated from WT or PS1 knockout (PS1 KO) mouse embryos were extracted as in 3C. VEGFR2 and VCTF1 were detected in cell extracts on WB with anti-VEGFR2 antibody as in 3B Left. β-actin: loading control. **Right:** HEK293T cells expressing VEGFR2-Myc were transfected with anti-PS1 siRNA or non-targeting control siRNA (NT siRNA) as in Methods and extracted as in 3B. VEGFR2-Myc and VCTF1-Myc were detected in cell extracts with anti-Myc antibody as in 3A. PS1 N-terminal fragment (PS1/NTF) was detected in cell extracts with anti-PS1 antibody (R222) [41]. β-actin: loading control. **(E), Left:** Extracts from WT and either WT/M146V- or WT/I213T-expressing pCECs were prepared as in 3C. VEGFR2 and VCTF1 were detected in cell extracts on WB with anti-VEGFR2 antibody as in 3B Left. β-actin: loading control. **Right:** Brain MVs were isolated from WT and either WT/M146V- or WT/I213T-expressing mice as in 1B and extracted as in 3C Right. VEGFR2 and VCTF1 were detected in MV extracts on WB with anti-VEGFR2 antibody as in 3B Left. β-actin: loading control. Each WB is representative of at least three independent experiments.

Fig 3C shows that treatment of WT mouse pCECs with RO accumulates VCTF1 (left). VCTF1 also accumulates in brain MVs prepared from RO-treated WT mice (right). Increased levels of VCTF1 are also found in pCECs prepared from PS1 knockout (PS1-/-) mouse embryos (Fig. 3D, left) and in HEK293T cells expressing reduced levels of PS1 (Fig. 3D, right) indicating that PS1 is required for the γ-secretase processing of VCTF1. γ-Secretase cleaves membrane-bound CTF1 substrates at the epsilon (ε) cleavage site usually located at the interface of the plasma membrane with the cytosol [31] and PS1 FAD mutations have been shown to decrease this γ-secretase-mediated cleavage [40–43]. To evaluate effects of PS1 FAD mutations on the γ-secretase processing of VEGFR2, we used WT pCECs and mice expressing FAD mutant PS1M146V or PS1I213T. Fig. 3E, left, shows that pCECs prepared from KI mice HTRZ for either one of the above FAD mutants accumulate VCTF1 compared to VCTF1 found in pCECs from WT mice. We found similar effects of PS1 FAD mutants on the accumulation of VCTF1 *<u>in vivo</u>* using brain MVs from HTRZ KI mice expressing either one of the above FAD variants compared to WT controls (Fig. 3E, right). Effects of additional PS1 FAD mutants on the γ-secretase–mediated cleavage of VEGFR2 were evaluated using the direct REC-LUC γ-secretase activity assay [57]. Our data show that five more PS1 FAD variants, E280A, A246E, G384A, E120K, and L166P transfected in HEK293T cell cultures, also reduce the γ-secretase cleavage of VEGFR2 (Suppl. Fig. 2). Together, these findings show that PS1 FAD mutations, RO GSI, and PS1 downregulation exert similar *<u>in vitro</u>* and *<u>in vivo</u>* effects on the γ-secretase-mediated processing of the VEGFR2-derived peptide VCTF1, reducing its catabolism and increasing its accumulation.

### 3. PS1 FAD mutants, RO GSI, and decreased PS1 all act to reduce the VEGF-A-induced dimerization of VEGFR2

To investigate how PS1 FAD mutations and RO influence the VEGF-A-induced angiogenesis, we examined their effects on VEGF-A-stimulated VEGFR2 dimerization, a critical angiogenic step regulating VEGFR2 autophosphorylation/activation and subsequent stimulation of downstream angiogenic signaling, EC functions, and angiogenesis [23,24]. To this end, we expressed a c-Myc-tagged VEGFR2 into HEK293T cells. Fig. 4A (three left lanes) shows that VEGF-A treatment stimulates the appearance of time-dependent and c-Myc-tag-positive high MW bands corresponding to the expected MW of VEGFR2 homodimers [24]. We also found that RO inhibitor blocks the VEGF-A-stimulated VEGFR2 dimerization (Fig. 4A, three right lanes). VEGF-A mainly stimulates formation of VEGFR2 homodimers [58] and we further verified homodimer formation using cells co-transfected with a VEGFR2-Venus N-terminus construct (VN, 1-172) and its complementary VEGFR2-Venus C-terminus construct [VC, 155-239] [59]. Transfected cells were then stimulated with VEGF-A in the presence or absence of RO and formation of VEGFR2 homodimers was estimated by measuring the green fluorescence produced by the VN-VC interaction. Our data (Suppl. Fig. 3A), show increased homodimerization in samples treated with VEGF-A only, but not in samples treated with vehicle, RO, or VEGF-A in the presence of RO. These data are in agreement with data in Fig 4A showing that RO inhibits VEGFR2 dimerization. Fig. 4B shows that MVs isolated from brains of WT mice treated with RO (see Methods), have decreased amounts of VEGF-A-induced VEGFR2 dimers compared to controls treated with VEGF-A only. Together, these data show that γ-secretase activity is required for VEGF-A-stimulated VEGFR2 dimerization *<u>in vitro</u>* and *<u>in vivo</u>*.

**Figure 4.**
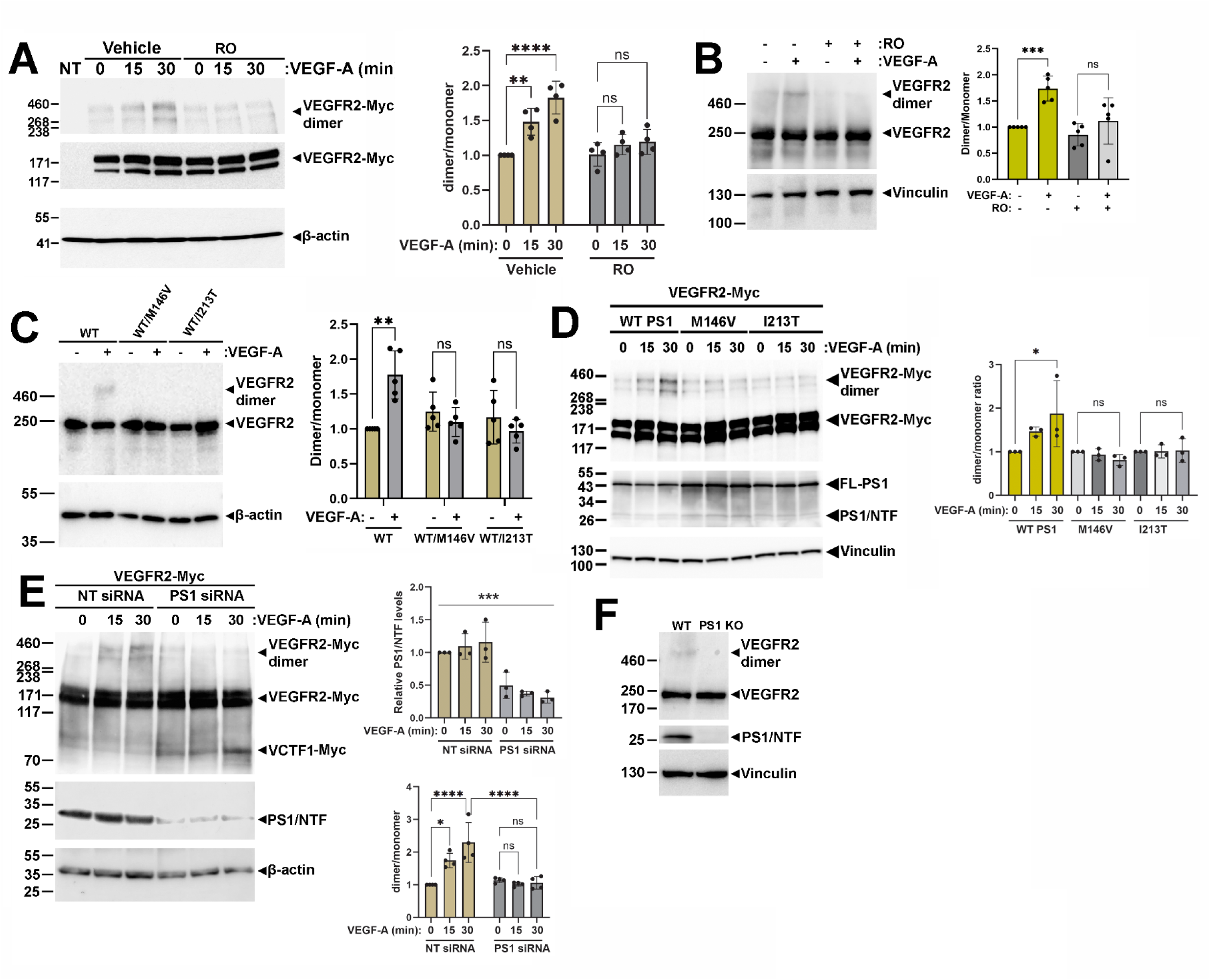
PS1 FAD mutants, GSIs and reduced PS1 suppress VEGF-A-induced VEGFR2 dimerization. **(A),** HEK293T cells were transfected with VEGFR2-Myc as in 3A and treated with vehicle or VEGF-A for the indicated times, in the presence or absence of RO as in 1F and extracted in SDS buffer. **Left:** VEGFR2 dimer and monomer were detected on WB with anti-Myc antibodies. β-actin: loading control. **Right:** Graph shows the fold change in the VEGFR2 dimer to monomer ratio. **(B),** WT mice were treated with vehicle or RO as in 1D. Mice were treated with VEGF-A, and brain MVs were isolated and extracted as in 1B. **Left:** VEGFR2 dimers and monomers were detected on WB with anti-VEGFR2 antibody D5B1. Vinculin: loading control. **Right:** Graph shows the fold change in the VEGFR2 dimer to monomer ratio. **(C),** WT and WT/M146V or WT/I213T mice were injected with either vehicle or VEGF-A via the carotid artery as in 1D, and brain MVs were prepared as in 1B and extracted as in 3C Right. **Left:** VEGFR2 dimers and monomers were detected on WB with anti-VEGFR2 antibody as in 4B Left. β-actin: loading control. **Right:** graph shows the fold change in the VEGFR2 dimer to monomer ratio. **(D),** HEK293T cells were co-transfected with VEGFR2-Myc as in 3A and either WT PS1 or PS1 mutant M146V or I213T in FCbAIGW vector as indicated in Figure. Cells were treated with VEGF-A as in 1G for the indicated times and extracted in SDS buffer. **Left:** VEGFR2 dimers and monomers were detected in cell extract on WB with anti-Myc antibody as in 4A. Full length PS1 (FL-PS1) and PS1/NTF were detected with R222 (middle). Vinculin: loading control. **Right:** Graph shows the fold change in the VEGFR2 dimer to monomer ratio. **(E),** HEK293 cells were co-transfected with VEGFR2-Myc and either non-targeting or anti-PS1 siRNA as in 3D. Cells were treated with vehicle (0 lanes) or VEGF-A as in 4D above for the indicated times and extracted in SDS buffer. **Left:** VEGFR2 dimers and monomers and VCTF1 were detected on WB with anti-Myc antibody as in 4A. PS1/NTF was detected in cell extracts with R222. β-actin: loading control. **Right:** graphs show fold change in PS1/NTF levels (upper) and fold change in the VEGFR2 dimer/monomer ratio (lower) following treatment with anti-PS1 siRNA. PS1 downregulation resulted in decreased VEGFR2 dimerization and increased VCTF1-Myc (upper panel). **(F),** Embryonic brain (E15.5) extract from WT or PS1 knockout (PS1 KO) mice were prepared as described [41]. Representative WB of extracts shows VEGFR2 dimers, detected with anti-VEGFR2 antibody D5B1, and PS1-NTF detected with R222 antibody. Vinculin: loading control. **A–E:** data are presented as mean ± SE from at least three independent experiments. Statistical analysis was performed by two-way ANOVA followed by Tukey post-hoc test. ns = not significant, *p<0.05, **p<0.01, ***p<0.001, ****p<0.0001.

To test effects of PS1 FAD mutants on the *<u>in vivo</u>* dimerization of VEGFR2, VEGF-A was injected into the carotid artery of WT mice and mice HTRZ for PS1 FAD mutants, and VEGFR2 dimers in brain MVs were detected as in Fig. 4B. Our data show that VEGF-A stimulates VEGFR2 dimerization in brain MVs of WT mice but not in brain MVs from mice expressing HTRZ PS1 FAD mutant M146V or I213T (Fig. 4C). We also found that cell cultures transfected with PS1 FAD mutant M146V or I213T have decreased levels of VEGF-A-induced VEGFR2 dimers compared to cultures expressing WT PS1 (Fig. 4D). Together, these data show that similar to RO, PS1 FAD mutants impair the VEGF-A-induced VEGFR2 dimerization both *<u>in vivo</u>* and *<u>in vitro</u>.* Five additional PS1 FAD variants—E280A, A246E, G384A, E120K, and L166P—were also tested via transfection in HEK293T cells and each exhibited a reduced effect on VEGF-A-induced VEGFR2 dimerization compared to WT (Suppl. Fig. 3B). Downregulation of PS1 also suppresses formation of VEGF-A-induced VEGFR2 dimers in cell cultures (Fig. 4E) and PS1 null (PS1- /-) embryonic mouse brains contain undetectable amounts of VEGFR2 dimers compared to WT controls (Fig. 4F) supporting that PS1 is required for VEGF-A-induced dimerization of VEGFR2. Furthermore, PS1 downregulation in cultured bEnd3 ECs, resulted in decreased levels of VEGF-A-stimulated pVEGFR2 and pERK1/2 (Suppl. Fig. 4A) and impaired EC tube formation (Suppl. Fig. 4B) indicating that reducing PS1 impairs angiogenic signaling and functions both of which take place downstream of the VEGFR2 dimerization [23,49]. Together, our data show that PS1 FAD mutants, RO GSIs or PS1 downregulation, all impair VEGFR2 dimerization. Combined with our findings that all three conditions above also reduce the γ-secretase cleavage of VCTF1 inducing its accumulation (Fig. 3), these data raise the possibility that increased levels of VCTF1 may impair VEGFR2 dimerization.

PS1 FAD mutants reduce γ-secretase processing of substrates [40–43] and in the HTRZ genotype can cause dominant negative (antimorphic) effects on the γ-secretase cleavage activity of the WT allele [43,60,61]. Accordingly, Suppl. Fig. 5A shows that pCECs HTRZ for FAD mutant PS1M146V or PS1I213T accumulate as much VCTF1 peptide as pCECs homozygous for the PS1 FAD mutants, indicating that HTRZ PS1FAD alleles have antimorphic effects on the γ-secretase processing of VCTF1. *<u>In vivo</u>* experiments also show that brain MVs from mice HTRZ for PS1 FAD mutants accumulate as much VCTF1 as that found in brain MVs from mice homozygous for FAD mutants (Suppl. Fig. 5B). Moreover, HTRZ PS1 FAD mutants completely abolish the VEGF-A-stimulated brain angiogenesis (Fig. 1A), angiogenic signaling such as activation of pVEGFR2 and pERK1/2 (Fig. 1E), EC tube formation (Fig. 2C), and VEGFR2 dimerization (Fig. 4C) despite presence of the WT PS1 allele. Together our data show that PS1 FAD mutants have dominant negative effects on the γ-secretase processing of VCTF1, abolishing the VEGF-A-induced VEGFR2 dimerization, signaling, EC functions and angiogenesis.

### 4. VCTF1 binds full-length VEGFR2 monomers, decreasing homodimerization

Our finding that PS1 FAD mutations, RO GSI, and PS1 reduction all increase VCTF1 accumulation and reduce VEGFR2 dimerization, led us to ask whether VCTF1 accumulation impairs VEGFR2 dimerization, thereby impairing VEGFR2 activation and attenuating downstream angiogenic signaling and EC functions. The apparent SDS-PAGE MW of VCTF1 (∼72 kDa) aligns well with the MW predicted from the 584 transmembrane and intracellular amino acids of VEGFR2 along with about 15 extracellular VEGFR2 residues adjacent to plasma membrane, totaling about 600 residues. We constructed a cDNA encoding the 600 amino acid sequence of VCTF1 tagged at C-terminus with an eight-residue c-Flag peptide (VCTF1-Flag, Suppl. Fig. 6) for detection purposes (see Methods). We introduced this cDNA into HEK293T cells co-expressing VEGFR2 tagged with c-Myc. Fig 5A (two left and two right lanes and graph below), shows that expression of the VCTF1-Flag inhibits the VEGF-A-stimulated dimerization of VEGFR2. In contrast, expression of a similarly tagged CTF1 peptide derived from an ADAM cleavage of EphB2 receptor (EphB2/CTF1, see Methods) and processed by γ-secretase [40] had no effect on the VEGF-A-stimulated dimerization of VEGFR2 (Fig. 5A, two left and two middle lanes). Moreover, Fig. 5B and graph below, show that increasing amounts of transfected VCTF1-Flag result in decreased levels of VEGF-A-induced VEGFR2 dimerization until plateau is reached at about a plasmid dosage of 0.4μg. Our data also indicated a half-maximal inhibitory concentration (IC_50_) of about 0.2μg (graph of Fig. 5B). Non-linear regression analysis showed a good fit to the curve (R² = 0.7410), indicating that VCTF1-Flag is an effective inhibitor of VEGFR2 dimerization. These data show that increased concentrations of VCTF1 result in decreased VEGF-A-induced VEGFR2 dimerization, supporting the hypothesis that PS1 FAD mutants, RO, and decreased PS1 may all impair VEGF-A-induced VEGFR2 dimerization by reducing the γ-secretase cleavage of VCTF1 increasing its concentration and enabling it to sequester VEGFR2 monomers. Additional evidence for an inverse relationship between VEGFR2-derived VCTF1 and VEGFR2 dimerization is shown in Fig 4E, where PS1 reduction (middle panel) of cell cultures transfected with VEGFR2-Myc monomer (upper panel) results in the accumulation of the VEGFR2-Myc-derived VCTF1-Myc (lower area of upper panel) with concomitant suppression of VEGFR2-Myc dimers (upper area of upper panel).

**Figure 5.**
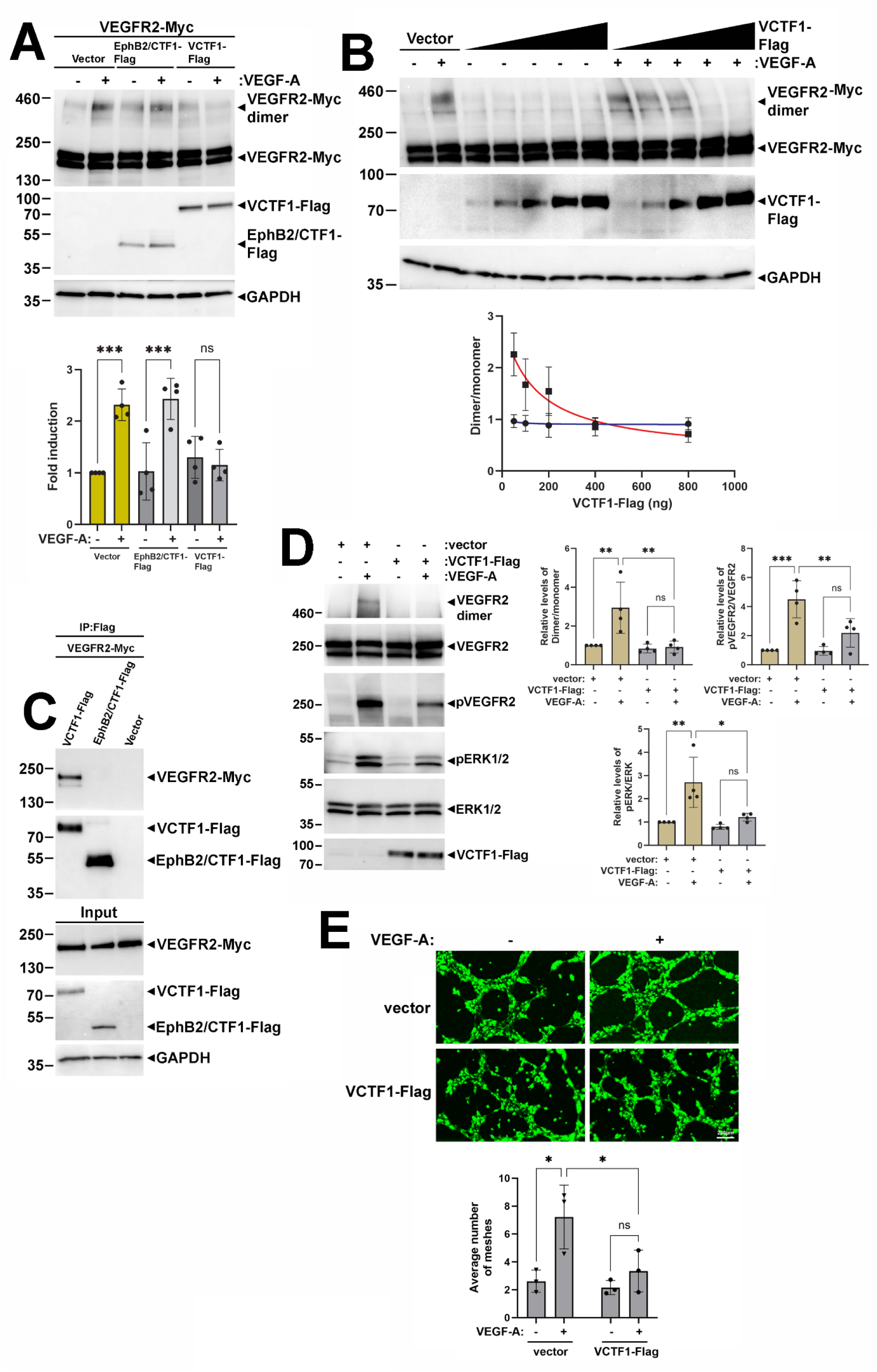
VCTF1 binds VEGFR2 monomers decreasing VEGF-A-induced VEGFR2 dimerization, downstream signaling, and angiogenic function. **(A),** HEK293T cells were co-transfected with vector expressing VEGFR2-Myc and plasmids expressing EphB2/CTF1-Flag, VCTF1-Flag, or vector alone. Cells were stimulated with vehicle (-) or VEGF-A (+) as in 1F and then extracted in SDS buffer. **Upper panel:** VEGFR2 monomers and dimers are detected on WBs using anti-VEGFR2 antibody OTI12C1. **Middle panel:** VCTF1-Flag and EphB2/CTF1-Flag expression was detected using anti-Flag antibody. **Lower panel:** GAPDH, loading control. Graph shows fold change of the VEGFR2 dimer to monomer ratio. **(B),** HEK293T cells were co-transfected with vector expressing VEGFR2-Myc and either vector alone or increasing amounts of vector expressing VCTF1-Flag (200ng, 400ng, 800ng, or 1000ng). Forty-eight hours post-transfection cells were stimulated with either vehicle or VEGF-A as in 1F for 20 min and extracted in SDS buffer. **Upper panel:** VEGFR2 dimers and monomers (VEGFR2-Myc) are detected in cell lysates on WBs using anti-VEGFR2 antibody OTI12C1. **Middle panel:** VCTF1-Flag is detected with anti-Flag antibody. **Lower panel:** GAPDH: loading control. **Bottom:** Non-linear regression analysis (inhibitor vs. response, three-parameter model) showed a good fit to the curve (R² = 0.7410), indicating that increase of VCTF1-Flag expression inhibits VEGF-A-induced VEGFR2 dimerization. The red line (squares) represents VEGF-A-treated cells, whereas the blue line (circles) represents vehicle-treated cells. **(C),** Cells described in 5A were lysed in Triton X-100 buffer as in Methods and lysates were IPed with anti-Flag antibody. **Upper panel:** VEGFR2-Myc co-IPed with VCTF1-Flag is detected on WB using anti-Myc antibody as in 3A. **Second panel:** IPed VCTF1-Flag and EphB2/CTF1-Flag are detected on WBs with anti-Flag antibody. **Third panel:** Input of VEGFR2-Myc is detected with anti-Myc antibody as in 3A. **Fourth panel:** Input VCTF1-Flag and EphB2/CTF1-Flag are detected with Flag antibody. GAPDH: loading control. **(D),** bEnd3 cells were transduced with lentiviral vector FCbAIGW expressing VCTF1-Flag or empty vector as in Methods. Cells were treated with vehicle (-) or VEGF-A (+) as in Suppl. 4A for 7 minutes and extracted in SDS buffer as in Methods. **Left:** VEGFR2 dimers, monomers, p-VEGFR2 (Tyr1175), p-ERK1/2, ERK1/2 and VCTF1-Flag are detected on WBs with specific antibodies. **Right:** Graphs show fold change of VEGFR2 dimer to monomer ratio or p-VEGFR2/VEGFR2 and p-ERK1/2/ERK1/2 protein ratios. **(E),** bend3 cells expressing either empty vector (FCbAIGW) or VCTF1-Flag as in 5D were seeded as in Methods and treated with vehicle (-) or VEGF-A (+) as in Suppl. 4A for 6 hours. **Upper:** Representative photomicrographs show tube-like (loop/mesh) structures. Fluorescent images (EGFP, green) are shown. Scale bar 200μm. **Lower:** Graph shows quantification of tube formation as average number of loops/meshes per field. For statistical analysis, two-way ANOVA followed by Tukey post-hoc test was performed. ns = not significant, *p<0.05, **p<0.01, ***p<0.001.

Since the TM and intracellular domains of VEGFR2 contribute to its dimerization [24,49], we asked whether VCTF1 might impair VEGFR2 dimerization by binding to VEGFR2 monomers, blocking formation of full-length homodimers. Using co-immunoprecipitation experiments, we obtained evidence that full-length monomers of VEGFR2 bind to VCTF1-Flag but not to EphB2-CTF1-Flag (Fig 5C). Furthermore, expression of VCTF1-Flag in bEnd3 brain ECs abrogates formation of VEGF-A-induced VEGFR2 dimers (Fig 5D, upper two panels) while also decreasing the VEGF-A-stimulated pVEGFR2 and pERK1/2 (Fig 5D, second to sixth panels from top). These data provide further support to the theory that suppression of VEGFR2 dimerization is a mechanism via which increased levels of VCTF1 impair downstream VEGF-A/VEGFR2-mediated angiogenic signaling and angiogenesis. The inhibitory effects of VCTF1-Flag on the VEGF-A-induced VEGFR2 dimerization and downstream signaling to pVEGFR2 and pERK1/2 indicate that VCTF1-Flag should also affect angiogenic functions of ECs. Indeed, Fig. 5E shows that transfection of VCTF1-Flag in bEnd3 ECs, strongly reduces VEGF-A-induced EC tube formation compared to vector transfected controls.

We also found that although VEGF-A stimulates VEGFR2 dimerization in brain MVs from WT mice, VEGF-A fails to stimulate VEGFR2 dimerization in brain MVs prepared from mice HTRZ for PS1 FAD mutants (Fig. 4C) which accumulate VCTF1 (Fig. 3E, right). Similarly, brain MVs from mice treated with RO accumulate VCTF1 (Figs. 3C, right) and show no increase in VEGFR2 dimerization in response to VEGF-A (4B). Moreover, downregulation of PS1 increases VCTF1 (Fig. 4E, upper panel marked VCTF1-Myc) while also reducing VEGF-A-induced VEGFR2 dimerization (Fig. 4E, upper panel, VEGFR2-Myc dimer). Together, our data support a model in which elevated brain VCTF1 increases its association with VEGFR2 monomers, thereby reducing the pool of monomers available for dimerization. This consequently diminishes both, VEGFR2 phosphorylation/activation and downstream angiogenic signaling and EC functions. Consistent with this interpretation, our findings show that both PS1 FAD mutants and RO inhibitors increase VCTF1 and suppress VEGFR2 signaling (Figs. 1D–E), trafficking (Figs. 1F, G), EC functions (Fig. 2), and brain angiogenesis (Figs. 1A and C).

### 5. PS1 FAD mutants block VEGF-A-induced neuronal survival and cognition under ischemic conditions

Brain ischemia is a major risk factor for neurodegeneration and dementia (13,20,21,62) and we used the MCAO mouse ischemia model [33] to explore effects of PS1 FAD mutants on VEGF-A-induced neuronal survival and cognition. Our data show that following MCAO-induced ischemia, neurons HTRZ for PS1 mutant M146V exhibit reduced survival on the side ipsilateral to the occlusion (Fig. 6A, right graph, bar 3 from left) compared to neuronal survival on the ipsilateral side of WT brains (Fig. 6A, right graph, bar 1, from left). We also found that the contralateral sides of HTRZ M146V and WT mouse brains had similar neuronal counts (Fig 6A, left graph) indicating that the PS1 FAD variant specifically increases neuronal vulnerability to ischemic stress. Consonant with the reduced neuronal survival caused by HTRZ mutant M146V, MCAO-treated mice expressing HTRZ PS1M146V showed a significantly greater impairment in cognition compared to MCAO-treated WT mice (Figs. 6B and C, compare yellow bar 3 from left with blue bar 5 from left). Combined with literature evidence that under ischemia, brain neuroprotection is mediated through the activation of VEGFR2 signaling by endogenous VEGF-A [22, 63–65], these findings raise the possibility that the HTRZ PS1 FAD mutant suppresses neuronal survival and cognition by targeting neuroprotective signaling of VEGFR2.

**Figure 6:**
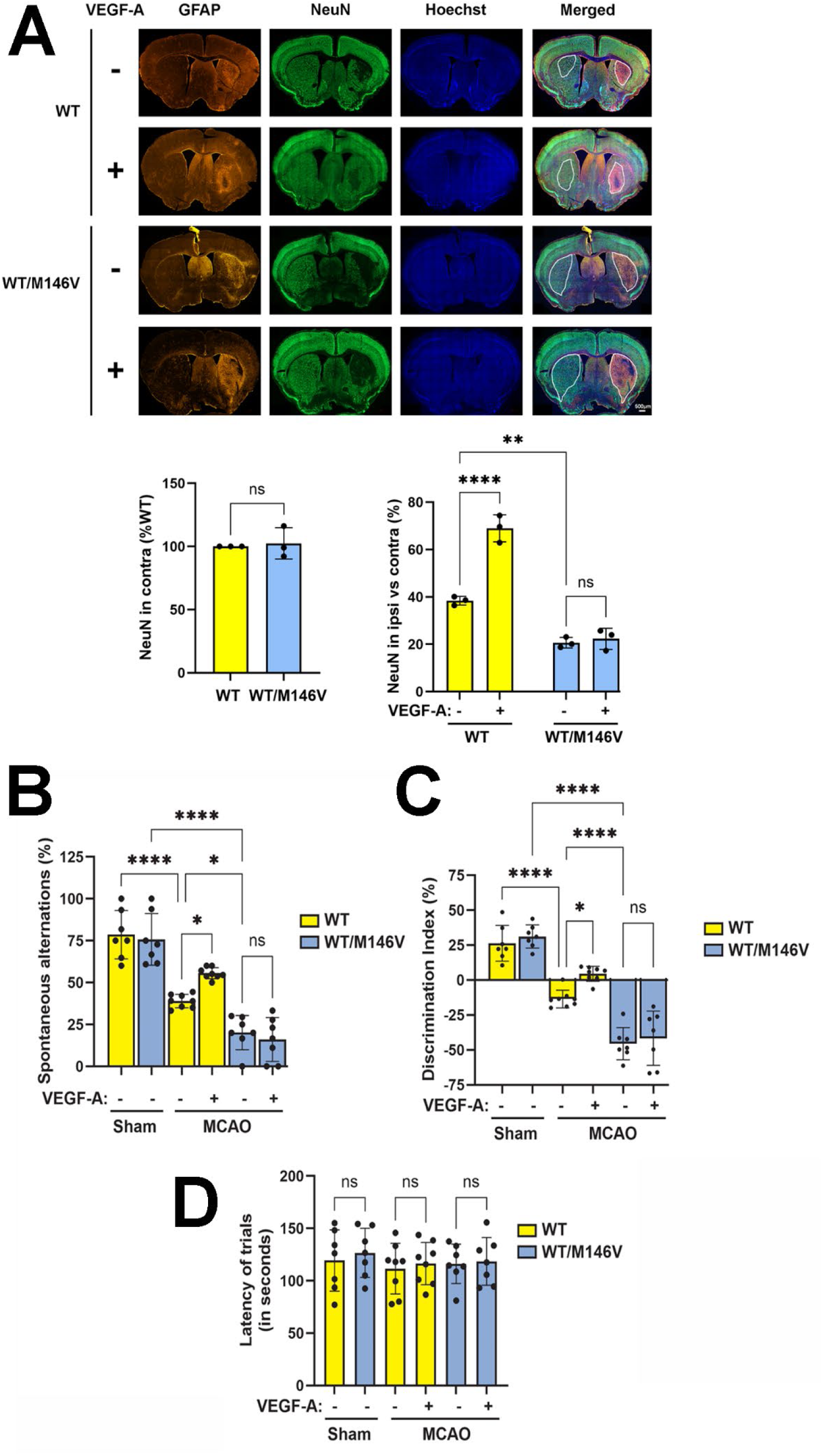
HTRZ PS1 FAD mutants sensitize neurons to Ischemia and abolish VEGF-A-induced neuroprotection and cognition. WT mice and mice HTRZ for PS1 WT/M146V mutant were subjected to MCAO as in Methods and then infused with vehicle or VEGF-A for 28 days as in 1A. **(A)**, brains were isolated and 30-μm-thick coronal sections were immunostained for NeuN (green), GFAP (red) and Hoechst (blue) as in Methods. **Upper:** Representative fluorescence photomicrographs (10X) show the lesion area. Scale bar: 500μm. **Lower left:** Graph represents NeuN-positive cell count (% of WT) in the contralateral side of each section. **Lower right:** Graph represents NeuN-positive cell count (%) in the lesion area of the ipsilateral side normalized to an identical size area in the contralateral side of each section as in Methods. **(B-D),** Mice were subjected to the following neurobehavioral tests as in Methods: **(B)** Y maze test, **(C)** Novel Object Recognition (NOR) test and **(D)** Rotarod test. Data represents Mean ± S.E. of at least seven mice per group, or as indicated in the dot plots. For statistical analyses, two-way ANOVA followed by Tukey post-hoc analysis was used. ns = not significant, *p<0.05, **p<0.01, ***p<0.001.

To further examine this hypothesis we asked whether the PS1 FAD mutant alters the effects of exogenous VEGF-A on neuronal survival and cognition. We noticed that, as expected [63–65], following MCAO ischemia, exogenous VEGF-A stimulates both neuronal survival (Fig. 6A, right graph, bars 1 and 2 from left) and memory capacity in WT mice (Figs. 6, B-D, yellow bars 3 and 4 from left). In contrast, our data here show that despite the presence of the WT PS1 variant, HTRZ PS1 FAD mutant completely abolishes both, the VEGF-A stimulated neuronal survival (Fig. 6A, right graph, WT/M146V blue bars 3 and 4 from left) and the VEGF-A stimulated memory recovery (Figs. 6B-D, WT/M146V, blue bars 5 and 6 from left) of ischemic mice. These data indicate that the HTRZ FAD mutant exerts dominant negative effects on neuroprotection. Although the mechanism(s) by which HTRZ PS1 FAD mutants abolish VEGF-A-induced neuronal survival and cognition are incompletely understood, a parsimonious explanation of our data is that the dominant negative effects exerted by the HTRZ PS1 mutant on the γ-secretase cleavage of VCTF1 (Suppl. Fig 5) and subsequent suppression of VEGFR2 dimerization (Fig. 4C) may be involved (see also Discussion).

### 6. VEGFR2 dimerization is reduced in human brain tissue from patients with FAD

We explored whether postmortem brain tissue from FAD patients expressing PS1 mutants have detectable molecular abnormalities relevant to the VEGFR2-mediated angiogenesis and neuroprotection. VEGFR2 dimerization is an early pivotal event that is required for both brain angiogenesis and neuroprotection [22,23,63,64] and Fig. 7A shows that human control brains have significantly higher amounts of VEGFR2 dimers than human brains of comparable age (Table 1) expressing PS1 FAD mutants. Despite our efforts, we detected neither peptide VCTF1 nor VCTF2, the product of the γ-secretase processing of VEGFR2, in human brain tissue. Literature shows that CTF1 γ-secretase substrates and CTF2 cleavage products are difficult to detect in postmortem brain tissue [66]. The VEGFR2 complex with endoglin is an important modulator of VEGFR2-mediated angiogenesis [39], and we discovered that this complex is significantly decreased in mouse brains expressing PS1 FAD mutants compared to WT controls (Fig. 1B). Fig. 7B shows that the VEGFR2/endoglin complex is also detectable in postmortem human brain tissue and significantly decreased in brains expressing PS1 FAD mutants compared to normal controls (Table 1). We also found that the cortical MV length density of control human brains is higher than the MV density of human brains expressing PS1 FAD mutants (Fig 7C; Table 2). Together, our findings show that human brain tissue expressing PS1 FAD mutants contains molecular signatures of impairments in VEGFR2 dimerization, VEGFR2/endoglin complex, and cerebral MV density

**Figure 7.**
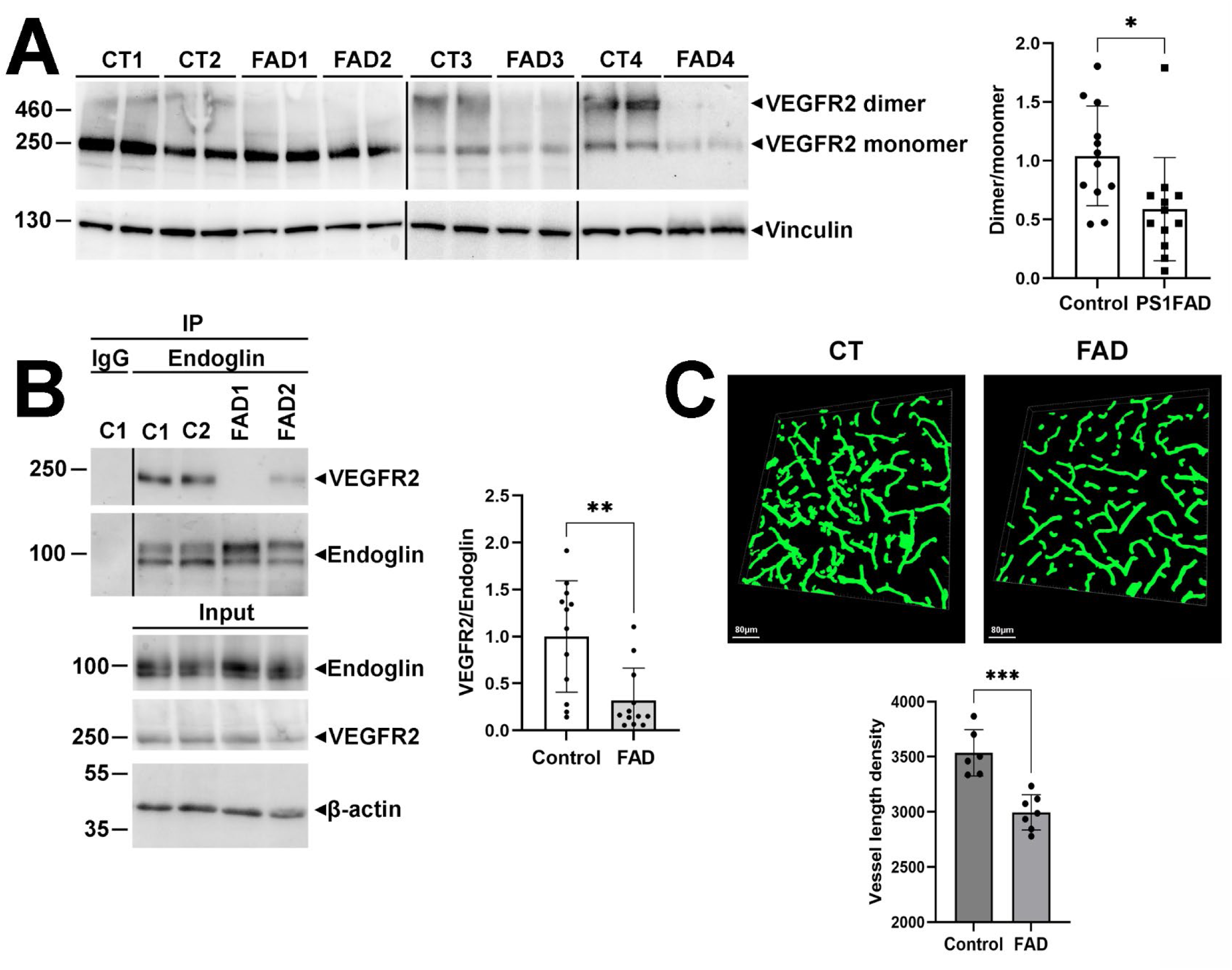
Decreased VEGFR2 dimers, VEGFR2/endoglin complexes, and brain vessel density in human PS1 FAD brains. **(A),** Brain tissue extracts were prepared as in Methods from twelve PS1 FAD patients each carrying a different PS1 mutation, and twelve non-demented controls described in Methods. **Left:** VEGFR2 dimers and monomers were detected in brain extracts on WBs using anti-VEGFR2 antibody D5B1. Representative gels with control (C1-4) or FAD samples (FAD1-4) expressing mutants P264L, A260V, N135S and P242H respectively are shown. Vinculin: loading control. **Right:** Graph shows the fold change in VEGFR2 dimer to monomer ratio of FAD and control samples. **(B),** Brain tissue extract from control and PS1 FAD patient brains described in 7A were prepared and IPed with anti-endoglin antibody (ab252345) or IgG as in Methods. **Upper panel:** VEGFR2 co-IPed with endoglin was detected on WBs using an anti-VEGFR2 antibody as in 7A. **Lower panel:** Input samples are shown. Representative gel with control samples (C1, C2) and FAD samples (FAD1, FAD2) expressing mutants A260V and P264L respectively is shown. β-actin: loading control. **Right:** Graph shows relative levels of VEGFR2 co-precipitated with endoglin. **(C),** Brain sections from control and PS1 FAD patients were prepared as in Methods and stained for Col IV as in 1A. **Upper:** Representative images show brain vessels in either PS1 FAD or control (CT) brain sections. Scale bar: 80μm. **Lower:** Graph shows total vessel length density in PS1 FAD and CT brains quantified with Imaris software as in 1A. **A-C**, bars represent Mean ± S.E. For statistical analysis, unpaired t-test was performed. *p < 0.05, **p<0.01 and ***p<0.001.

**Table 1:** Human brain tissue used in Fig. 7A and 7B.

| WB (7A), IP (7B) | Mutant | Sex | Age |
| --- | --- | --- | --- |
| PS1 FAD | I143T | F | 36 |
|  | I143T | F | 38 |
|  | N135S | M | 43 |
|  | G206A | M | 64 |
|  | G206A | M | 70 |
|  | P242H | F | 63 |
|  | P264L | F | 57 |
|  | A260V | F | 65 |
|  | H163R | M | 51 |
|  | I229F | F | 39 |
|  | S170F | M | 40 |
|  | G206V | M | 40 |
| Controls |  | F | 63 |
|  |  | F | 56 |
|  |  | M | 48 |
|  |  | F | 70 |
|  |  | F | 54 |
|  |  | M | 70 |
|  |  | F | 24 |
|  |  | M | 37 |
|  |  | M | 67 |
|  |  | F | 55 |
|  |  | F | 79 |
|  |  | M | 52 |
|  |  | F | 66 |
|  |  | F | 67 |

**Table 2:** Human brain tissue used in Fig. 7C.

| IHC (7C) | Mutant | Sex | Age |
| --- | --- | --- | --- |
| PS1 FAD | A398V | F | 65 |
|  | V272A | F | 40 |
|  | G206A | F | 77 |
|  | A79V | F | 70 |
|  | M146L | M | 38 |
|  | H163R | M | 51 |
|  | I249L | M | 72 |
| Controls |  | F | 42 |
|  |  | F | 66 |
|  |  | F | 70 |
|  |  | F | 78 |
|  |  | M | 48 |
|  |  | M | 74 |

## Discussion

Brain angiogenesis and neovascularization are tightly regulated processes playing critical roles in both development and post-injury repair of adult vasculature. VEGF-A stimulates brain angiogenesis and neovascularization by binding VEGFR2 (also known as KDR or Flk1), the main mediator of VEGF-A-induced angiogenesis, thus activating VEGFR2-mediated angiogenesis cascades [22,23]. Importantly, VEGFR2 signaling also contributes to neuronal survival after insult through neurotrophic and neuroprotective actions [22, 63–65]. Here we describe a γ-secretase-dependent mechanism that regulates VEGF-A-induced and VEGFR2-mediated brain angiogenesis. Specifically, our data show that GSI RO suppresses VEGF-A-induced mouse brain angiogenesis and impairs VEGFR2-mediated angiogenic functions such as VEGF-A-induced VEGFR2 dimerization, phosphorylation/activation and trafficking. Moreover, RO suppresses activation of kinases important to angiogenesis and neuroprotection such as ERK1/2 [23,65], and reduces angiogenic functions of ECs including sprouting, migration and tube formation. We also found that PS1 FAD mutants have similar negative effects on VEGF-A-stimulated brain angiogenesis to those caused by the RO inhibitor, suppressing VEGF-A-induced VEGFR2 dimerization, activation, signaling, and EC functions and angiogenesis. The strong similarities between RO-induced impairments of brain angiogenesis and VEGFR2 signaling and functions and those observed in PS1 FAD mutants suggest that PS1 FAD mutations act as GSIs, severely impairing the γ-secretase cleavage activity of PS1 [40–43]. Together, our findings indicate that γ-secretase activity regulates VEGF-A/VEGFR2-mediated brain angiogenesis, and that both RO and PS1 FAD mutants reduce VEGFR2-mediated angiogenesis by suppressing the VEGFR2 dimerization and disrupting the subsequent downstream signaling of the VEGFR2 [22,23].

VEGF-A stimulates an ADAM-17 cleavage and shedding of the extracellular domain of VEGFR2 [29], and we show here that the remaining membrane bound VEGFR2-derived peptide VCTF1 is further processed by γ-secretase. We also found that cell cultures and brain MVs from mice expressing PS1 FAD mutants accumulate the γ-secretase substrate VCTF1. As expected, VCTF1 also accumulates in EC cultures treated with RO and in brain MVs from RO-treated WT mice. Moreover, VCTF1 accumulates in PS1 null (PS1-/-) EC cultures and in HEK cell cultures expressing reduced PS1. These data show that VCTF1 accumulates under conditions of decreased γ-secretase activity and that PS1 FAD mutations stimulate accumulation of VCTF1 by decreasing the γ-secretase activity of PS1. VEGF-A-induced VEGFR2 dimerization is a key early step in angiogenesis, controlling VEGFR2 phosphorylation/activation and downstream angiogenic signaling, EC functions, and angiogenesis [22–24], (also Fig. 8). We found that in addition to stimulating VCTF1 accumulation, PS1 FAD mutants, RO, and PS1 reduction, all impair VEGF-A-induced VEGFR2 dimerization in EC cultures and brain tissue (Fig. 4). In agreement with our findings that PS1 downregulation causes VCTF1 accumulation decreasing VEGFR2 dimerization, PS1 reduction also reduces downstream phosphorylation/activation of both VEGFR2 and ERK1/2 (Suppl. Fig. 4A) and suppresses VEGF-A-induced EC tube formation (Suppl. Fig. 4B). These effects are similar to those caused by GSIs and PS1 FAD mutants both of which accumulate VCTF1. Together, these findings show that γ-secretase activity is required for efficient VEGF-A-induced VEGFR2 dimerization, activation, angiogenic signaling and EC functions. Our data also show that although PS2 protein is expressed in all biological systems used here, it does not compensate for deficits caused on VCTF1 processing by PS1 reduction and ensuing decreases in VEGFR2 dimerization, signaling, or EC functions. These data are consistent with evidence that certain γ-secretase substrates, including N-cadherin and ephrinB, are processed exclusively by PS1 [31,32] and that PS1 and PS2 function at different subcellular localizations with PS1 being mainly responsible for γ-secretase activity at the plasma membrane [56].

**Figure. 8:**
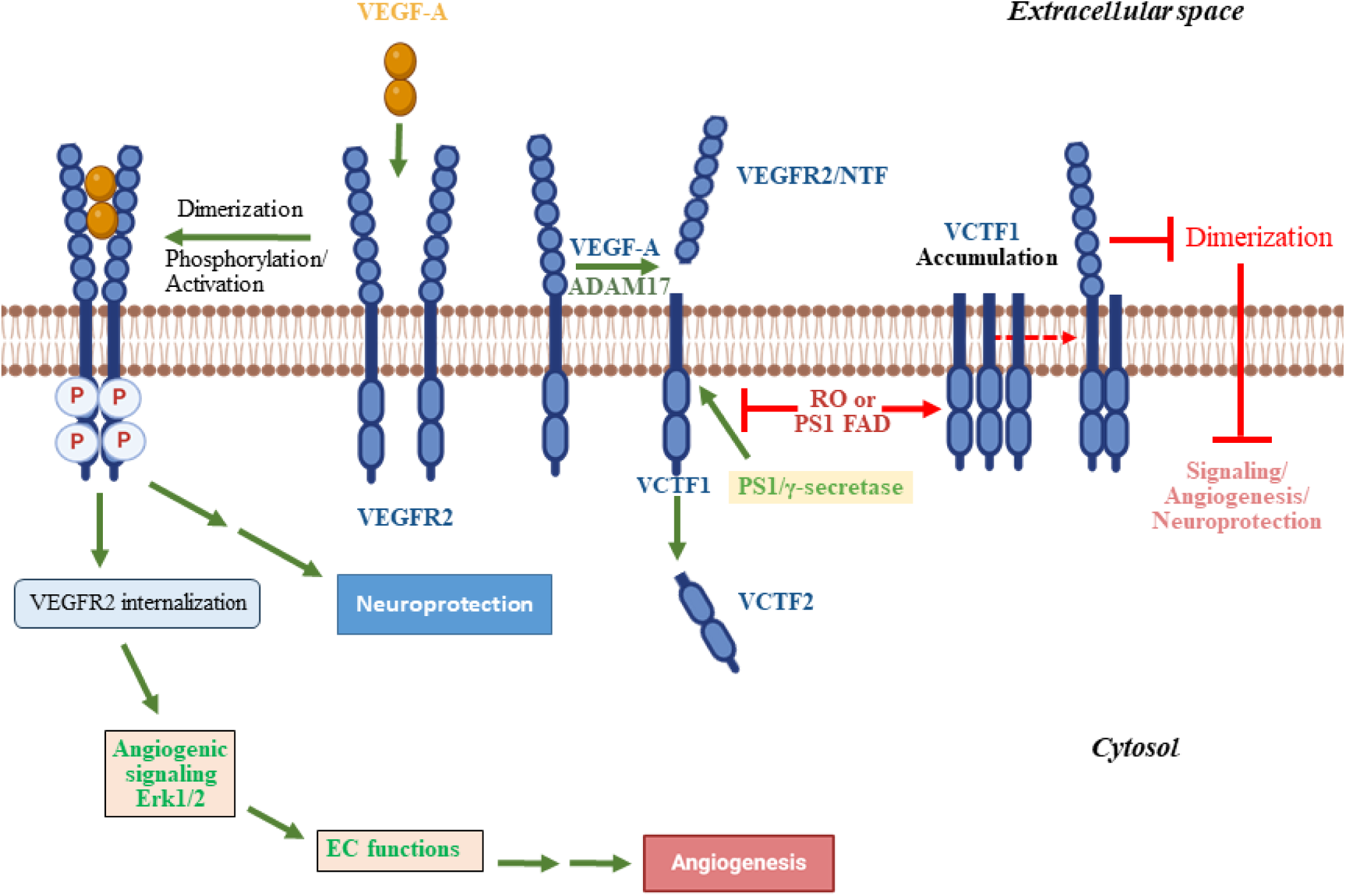
PS1 FAD mutants or GSIs reduce VCTF1 cleavage increasing its accumulation. VEGFR2 is processed by ADAM metalloproteinases (MPs) near the extracellular face of the plasma membrane, producing an ectodomain peptide, termed VEGFR2/N-terminal fragment (NTF) shed to medium, and a membrane-bound peptide termed VCTF1. PS1 FAD mutants or RO reduce the cleavage of VCTF1, leading to its accumulation on the membrane. Accumulated VCTF1 interacts with VEGFR2 monomers preventing VEGF-A-induced VEGFR2 homodimerization and impairing angiogenic signaling and function. At left is an abbreviated diagram of the activated VEGFR2 pathway to both angiogenesis and neuroprotection.

Our finding that reduction of γ-secretase activity results in VCTF1 accumulation and suppression of VEGFR2 dimerization, angiogenic signaling, and EC functions, prompted us to explore further potential VCTF1 roles on the VEGF-A-induced dimerization of VEGFR2 and its subsequent downstream signaling. To this end, we constructed a cDNA encoding the amino acid sequence of peptide VCTF1 tagged with c-Flag (VCTF1-Flag, Suppl. Fig. 6). Our data show that (1) VCTF1-Flag impairs VEGFR2 dimerization (Fig. 5A), and increasing concentrations of VCTF1-Flag progressively decrease VEGFR2 dimerization (Fig. 5B). (2) VCTF1-Flag is an effective inhibitor of VEGFR2 dimerization as indicated by non-linear regression (Fig. 5B). (3) Expression of VCTF1-Flag in ECs blocks VEGF-A-induced VEGFR2 dimerization and reduces pVEGFR2, pERK1/2 (Fig. 5, D), and EC tube formation (Fig. 5E), indicating that the presence of VCTF1 impairs VEGFR2 dimerization, activation and downstream angiogenic signaling and EC functions. (4) Mouse brain MVs that accumulate VCTF1 in response to either RO treatment or expression of PS1 FAD mutants (Figs. 3, C and E respectively) also show decreased VEGF-A-induced VEGFR2 dimerization (Figs 4, B and C respectively) and reduced pVEGFR2 and pERK1/2 formation (Figs. 1, D and E respectively). Moreover, both mice treated with RO and those expressing PS1 FAD mutants show decreased VEGF-A-induced angiogenesis (Figs. 1A, C). (5) An inverse relationship between VEGFR2-derived VCTF1 and VEGFR2 dimerization is depicted in Fig 4E, where reduction of PS1 in HEK293T cell cultures transfected with VEGFR2-Myc results in increased accumulation of the VEGFR2-Myc-derived VCTF1-Myc and the concomitant suppression of VEGFR2-Myc dimerization. Combined, our data show that increased cellular levels of VCTF1 impair VEGF-A-induced VEGFR2 dimerization, leading to decreased VEGFR2 activation, signaling, EC functions, and angiogenesis.

Our finding that full-length VEGFR2 binds VCTF1 (Fig. 5C) supports a mechanism by which increased accumulation of cellular VCTF1 binds increased amounts of full-length VEGFR2 monomers reducing their availability for homodimerization, consistent with our data that increased amounts of VCTF1 result in decreased VEGFR2 dimerization (Figs. 4E and 5, A, B and D). Moreover, PS1 FAD mutations, RO treatment, and PS1 reduction, all decrease γ-secretase processing of VCTF1, increasing its accumulation (Fig 3,C-E) and suppressing VEGFR2 dimerization (Fig. 4), an outcome that reduces downstream VEGFR2 activation, signaling, EC functions, and angiogenesis [23], (also Fig 8). Taking advantage of the observation that expression of exogenous PS1 or PS1 FAD variants in cell cultures reliably indicates mutation effects on VEGFR2 dimerization, we assessed the impact of five additional PS1 FAD variants on VEGFR2 dimerization. We found that expression of each one of seven distinct PS1 FAD mutants in cell cultures caused a significant reduction in the VEGF-A-induced VEGFR2 dimerization (Fig. 4D and Suppl. Fig. 3B), an outcome indicating that many PS1 FAD mutants impair VEGFR2 dimerization. Together, our data show that accumulation of the γ-secretase substrate VCTF1 decreases VEGFR2 dimerization reducing its phosphorylation/activation and downstream angiogenic signaling and EC functions, and suppressing angiogenesis. These observations raise the possibility that interventions that reduce brain VCTF1 may increase brain angiogenesis and consequently blood flow benefiting FAD patients.

Brain vascular damage can induce ischemia, a harmful cycle of energy failure, excitotoxicity, and oxidative stress culminating in brain infarction and neuronal death. The brain reacts to ischemia by increasing production of VEGF factors that activate VEGFR2 stimulating both angiogenesis and neuroprotective signaling [22,63,64]. We noticed that MCAO treatment of mice HTRZ for mutant PS1M146V results in significantly greater neuronal loss compared to neuronal loss in MCAO-treated WT mice (Fig. 6A, right graph, bars 1 and 3 from left). Moreover, MCAO treatment of mice HTRZ for PS1M146V exhibit significantly impaired cognitive performance relative to WT mice subjected to the same MCAO treatment (Figs 6, B,C, bars 3 and 5 from left). Since under ischemia, neuroprotection is mediated by endogenous VEGF-induced activation of VEGFR2 [22,63,64], our data raise the possibility that mutant PS1M146V interferes with functions of the VEGF-A/VEGFR2 neuroprotective system, rendering brain neurons more vulnerable to ischemic insult.

To explore further potential effects of mutant PS1M146V on the neuroprotective functions of the VEGF-A/VEGFR2 system, we tested effects of exogenous VEGF-A on the neuroprotection of WT mouse brains and mouse brains expressing mutant PS1M146V. We found that, as expected [22,64], following MCAO, treatment of WT mice with VEGF-A increased both neuronal survival and cognition. In contrast, VEGF-A treatment of mice HTRZ for mutant M146V increases neither neuronal survival nor cognition (see results, section 5 and Figs. 6, A-D). Our data show that mutant PS1M146V causes accumulation of VCTF1 (Fig. 3,E) and suppression of VEGF-A-induced VEGFR2 dimerization (Fig. 4C) and activation (Fig. 1E). Since VEGFR2 dimerization is required for VEGFR2-mediated neuroprotection [22, 63–65], a reasonable explanation of our data is that VCTF1 accumulation decreases VEGF-A-induced neuroprotection by suppressing VEGFR2 dimerization. Together, these findings support the theory that PS1 FAD mutants impair neuroprotection and cognition by accumulating VCTF1 and suppressing VEGFR2 dimerization and signaling, functions that are essential for VEGFR2-mediated neuroprotection after ischemic insult [22,63,64].

Demonstrating the relevance of our findings to human AD patients, we also found that human brain tissue expressing PS1 FAD mutants exhibit molecular signatures indicative of impaired VEGFR2 dimerization, VEGFR2/endoglin complex formation, and angiogenesis (Fig. 7). These effects mirror findings in brains of mice expressing PS1 FAD mutants. Combined, our data suggest a pathway through which FAD mutants promote dementia by increasing VCTF1 and suppressing brain angiogenesis, neuroprotection and cognition. In addition, our findings suggest that therapeutic strategies aimed at enhancing the metabolism and/or clearance of VCTF1 and promoting VEGFR2 dimerization may be beneficial to PS FAD patients. Finally, our data indicate that the γ-secretase-dependent clearance of VCTF1 is a *p*reviously unrecognized regulatory checkpoint in the biology of VEGFR2. By impairing VEGFR2 dimerization and suppressing downstream angiogenesis and neuroprotection, VCTF1 accumulation may also have pathogenic implications in additional diseases where VEGF/VEGFR2 signaling plays important roles, including stroke and cancer.

## Data availability

Data used and/or analyzed for this study are either included in article (and its Supplementary Information) or are available from corresponding authors on request.

## Ethics statement for the use of human tissues and animals

### Human tissue

The use of human post-mortem tissue complied with institutional ethical guidelines and the principles of the World Medical Association Declaration of Helsinki.

### Animals

The experimental procedures were conducted in accordance with NIH guidelines for animal research and were approved by the Institutional Animal Care and Use Committee (IACUC) at the Icahn School of Medicine at Mount Sinai.

## Author Contributions

All lab members participated in discussing Ideas and planning experiments to test questions.

All authors contributed to the interpretation of outcomes.

N.K.R., A.G. approved the experiments. A.G., N.K.R. and R.P. supervised the work.

R.P., A.Z., P.D. and E.L. carried out the experiments.

N.K.R. and A.G. wrote the manuscript and A.G and R.P. designed the figures.

A.G. and R.P. designed the computational framework and analyzed the data.

G.C. analyzed tube formation data.

P.H. provided consultation on the processing of brain tissue and quantification of IHC data.

## Materials and Methods

### Animals

All experimental procedures followed the NIH Guide for Care and Use of Laboratory Animals approved by the Institutional Animal Care and Use Committee (IACUC) of the Icahn School of Medicine at Mount Sinai. Mice were maintained under standardized housing conditions, including a 12-hour light/dark cycle, controlled room temperature (RT; 23 °C), and unrestricted access to food and water. Experimental cohorts included WT and KI mice expressing FAD mutant M146V (35) or I213T [36], and PS1-Knockout mice [67] previously characterized [33]. For experiments 3-4 month-old male C57BL/6 mice were used.

### Materials and antibodies

Rabbit monoclonal anti-VEGFR2 (ab39256), anti-Collagen IV (Col IV) (ab6586) and anti-endoglin (CD-105) (ab252345) were from Abcam (Cambridge, MA), anti-VEGFR2 (D5B1), anti-p-VEGFR2 (Tyr1175) (D5B11), anti-ERK1/2 (137F5), anti-p-ERK1/2 (D13.14.4E), anti-β- actin (13E5) and anti-vinculin (E1E9V) from Cell Signaling Technologies (Beverly, MA). Rabbit polyclonal anti-Col IV (AB756P), anti-NeuN (ABN78) from Millipore Sigma, anti-Myc tag (2272) from Cell Signaling Technologies (Beverly, MA), anti-p-VEGFR2 (Tyr1054/Tyr1059) (cat#44-104-7G) from Fisher Scientific and anti-PS1 antibody (R222) has been previously described [41]. Mouse monoclonal anti-Flag tag (M2; F1804) was from Millipore Sigma, anti-GAPDH (2118S) from Cell Signaling Technologies (Beverly, MA), anti-VEGFR2 (OTI12C1) from Origene, anti-endoglin (CD-105; NBP2-22122) and anti-LAMP2 (NBP2-22217) from Novus Biologicals, Inc, anti-Rab5 (D-11) and anti-Rab7 (B-3) from Santa Cruz Biotechnology, Inc. Chicken polyclonal anti-GFAP (ab4674) was from Abcam. Peroxidase-conjugated AffiniPure goat anti-mouse IgG light chain-specific were from Jackson Immunoresearch Laboratories Inc. Goat anti-Rabbit IgG (H+L) Alexa Fluor 488, goat anti-Mouse IgG Alexa Fluor 568 and goat anti-Chicken IgY (H+L) Alexa Fluor Plus 555 were from Invitrogen. Anti-Rabbit IgG (H+L), peroxidase-labeled, was from Clinical Diagnostics Inc.

Recombinant mouse VEGF164 (493-MV) was from R&D Systems. EC medium without FBS (MCDB131) was from VEC Technologies. ECGS Bovine supplement (CB40006B), EGF Mouse supplement (CB40010), DMSO (BP231100) and protease inhibitor cocktail (501933279) were from Fisher Scientific. Dynabeads protein A or G (10001D and 10003D respectively) were from Invitrogen. Lactacystin was from VWR International (Avantor; ABCA_AB141411). γ-Secretase inhibitor RO4929097 (S1575), Compound E (S0058), SU5416 (Semaxanib; S2845) and PEG300 (S6704) were from Selleck Chemicals LLC. ADAM17 inhibitor (clone D1(A12)) was from Millipore Sigma. Cytodex® microcarrier (MC) beads, fibrinogen (type I-S from bovine plasma), aprotinin, thrombin (from bovine) ECGS (354006), EGF (354001) and Hoechst 33258 were from Sigma-Aldrich Inc. Lentivirus concentrator (NC9833735) was from Fisher Scientific LLC.

### Middle cerebral artery occlusion (MCAO)

Focal cerebral ischemia was induced in adult male mice 3-4 months (25–30 gr) as described [68,33]. Successful MCAO was monitored as we described [33].

### Administration of VEGF-A, RO and SU in mice

One time carotid artery injections were performed to deliver RO4929097 or VEGF-A into cerebral circulation, following previously described procedures [69]. Briefly, adult male mice 3-4 months old (25–30 g) were anesthetized with isoflurane, and a midline cervical incision was made to expose the left common carotid artery (CCA), external carotid artery (ECA), and internal carotid artery (ICA). A catheter connected to a syringe prefilled with the treatment solution was positioned for infusion. A small incision was made in the CCA, and the catheter tip was carefully advanced toward the ICA. RO4929097 or VEGF-A was slowly injected into the ICA to ensure uniform distribution to the cerebral vasculature [69]. Following injections, the catheter was withdrawn, and incision site was ligated. Brains were harvested at the indicated time points (as detailed in figure legends), and MVs were isolated as described below. RO4929097 was prepared as a 10 mg/mL stock in 2% DMSO, 30% PEG300, and 1% Tween-80 in water. Immediately prior to injection, the stock was diluted in sterile 0.9% saline to working concentrations of 3 mg/mL (tail vein; 50 µL; 5 mg/kg for a 30 g mouse) or 1 mg/mL (carotid artery; 30 µL; 1 mg/kg for a 30 g mouse). SU5416 (semaxanib; Selleck Chemicals) was prepared as a 10 mg/mL stock in a vehicle containing 5% DMSO, 40% PEG300, 5% Tween-80, and 50% ddH₂O. Mice received SU5416 at 20 mg/kg from 10 mg/mL stock. Delivery of RO4929097 and SU4516 in ischemic mice was done by tail vein and intraperitoneal (IP) injections respectively as described [70,71].

### Mini osmotic pump implantation for carotid artery delivery of VEGF-A

Experiments were conducted in 3–4-month-old mice (25–30 g). WT and PS1 M146V KI mice were used for VEGF-A infusion through a mini-osmotic pump (Alzet, DURECT Corporation) as described previously [37]. Briefly, under isoflurane anesthesia, a midline incision was made in the neck to expose the left CCA, ECA and ICA. A custom catheter (MJC-28, SAI Technologies) was introduced into the ECA and advanced up to the carotid bifurcation so that the catheter opening faced the ICA, ensuring infusion toward the cerebral circulation. The catheter was connected to a mini-osmotic pump prefilled with either VEGF-A solution or vehicle (0.2% BSA in PBS).

The osmotic pump model was selected according to the desired infusion duration. Model 1002 (100 µl reservoir, 0.25 µl/h flow rate) was used for 2-week infusions, and Model 1004 (100 µl reservoir, 0.11 µl/h flow rate) for 4-week infusions. Pumps were implanted subcutaneously on the dorsal surface to deliver the contents continuously over the specified period. Following surgery, animals were monitored until full recovery and maintained under standard postoperative care conditions.

### Image acquisition and analysis: (A) Brain

Images were acquired on a Zeiss LSM980 microscope (Carl Zeiss Microscopy GmbH, Jena, Germany), using Zen Blue software (3.7.97), equipped with multi- alkali and GaAsP Pmt detectors (Hamamatsu Photonics, Shizuoka, Japan) in confocal mode. Imaging was performed using a 20x/0.8 Plan-Apochromat lens (Carl Zeiss Microscopy GmbH, Jena, Germany). (1), Measurement of Col lV in the absence of MCAO: Microscope parameters: Optical zoom – 1.2; Image size – 512x512 pixels; Pixel size – 0.691μm/pixel; Image area – 354x354μm; Bit depth – 8 bits; Pixel dwell time – 0.85μs/pixel; Scanning – bi-directional; Scan Mode – Frame; Lasers – 405nm Diode and Argon 488nm for DAPI and Alexa488, respectively; Dichroics – MBS 405 and MBS488/561 double dichroic (Chroma Technology, Bellows Falls, VT, USA), Emission windows – 416-491nm and 500-553nm for DAPI and Alexa488, respectively; Detectors – Multi-alkali Pmt and GaAsP Pmt for DAPI and Alexa488, respectively; Airy unit – 1; Pinhole size – 24μm; Z-step size – 1.5μm/step; Z-thickness – 40μm (27 z-slices). (2), For measurement of NeuN following MCAO: Optical zoom – 1; Pixel size – 0.830μm/pixel; Scanning – Tiles; Bit depth – 8 bits; Pixel dwell time – 2.05μs/pixel; Scanning – bi-directional; Scan Mode – Frame; Lasers – 405nm Diode, Argon 488nm and Diode 561nm for DAPI, Alexa488 and Alexa555, respectively; Dichroics – MBS 405 and MBS488/561 double dichroic (Chroma Technology, Bellows Falls, VT, USA), Emission windows – 424-489nm, 500-561nm and 575-698nm for DAPI, Alexa488 and Alexa555, respectively; Detectors – Multi-alkali Pmt, GaAsP Pmt and Multi-alkali PMT2 for DAPI, Alexa488 and Alexa555, respectively; Airy unit – 1; Pinhole size – 34μm; Z-step size – 1.5μm/step; Z-thickness – 30-35μm (21-24 z-slices). (3), For measurement of Col lV following MCAO: Acquisition was performed as in 2; Z-thickness – 39μm (27 z-slices). (4), For measurement of Col lV in human brain tissue: Optical zoom – 0.7; Pixel size – 1.18μm/pixel; Scanning – Tiles; Bit depth – 8 bits; Pixel dwell time – 1.54μs/pixel; Scanning – bi-directional; Scan Mode – Frame; Lasers – 405nm Diode and Argon 488nm for DAPI and Alexa488, respectively; Dichroics – MBS 405 and MBS488/561 double dichroic (Chroma Technology, Bellows Falls, VT, USA), Emission windows – 424-489nm and 500-561nm for DAPI and Alexa488, respectively; Detectors – Multi-alkali Pmt and GaAsP Pmt for DAPI and Alexa488, respectively; Airy unit – 1; Pinhole size – 28 μm; Z-step size – 1.5μm/step; Z-thickness – 30μm (21 z-slices). Raw images were converted into Imaris-compatible files using the Imaris File Converter software. Preprocessing steps included background subtraction and Gaussian filtering to reduce noise and enhance vascular signal clarity. The Filament Tracer module in Imaris was used to quantify vessel parameters with a minimal ratio of branch length to trunk ratio of 1.85. Vessel length was calculated from the total path length of filamentous structures representing the vasculature. The vessel length density was determined by normalizing the total vessel length to the corresponding imaged volume. Only MVs with diameter < 50μm were measured.

For neuronal count in lesioned area neurons were stained with anti-NeuN antibody and lesion area was identified with anti-GFAP staining. The number of NeuN-immunolabelled neurons in lesion area and contralateral area were counted in every 10^th^ section using the Imaris software. Counting was performed automatically after manually creating surfaces that include the neurons in the lesion area to be counted and then counting the number of objects in the corresponding surface. Under low magnification, the boundary of lesion area was identified with GFAP staining and the boundary contour in lesion area was drawn using the ImageJ software. The same drawn area was used in the contralateral lateral area of the section.

### (B) pCEC

Images were acquired with LAS X software (3.5.7.23225) on a Leica SP8 – STED 3x confocal microscope (Leica Microsystems, Wetzlar, Germany) equipped with multi-alkali Pmts and HyD detectors. Imaging was performed using a HC-PL-APO-CS2 100x/1.4NA oil lens (Leica Microsystems). Microscope parameters : Optical zoom – 1; Image size – 1024x1024 pixels; Pixel size – 0.11μm/pixel; Image area – 116.25x116.25μm; Bit depth – 16 bits; Pixel dwell time – 0.4μs/pixel; Scanning – uni-directional; Scan Mode – Line; Lasers – 405nm Diode, WLL 499nm and WLL 553nm DAPI, Alexa488 and Alexa 555, respectively; Dichroics – TD 405 and TD 488/561/633 triple dichroic, Emission windows – 410-504nm, 504-548nm and 558-725nm for DAPI, Alexa488 and Alexa555, respectively; Detectors – Multi-alkali Pmt and HyD detectors for DAPI, Alexa488 and Alexa555, respectively; Pinhole size – 151μm; Z-step size – 0.1μm/step; Z-thickness – 2.49μm (26 z-slices). Image analysis was conducted using Imaris software to quantify colocalization between VEGFR2 and endosomal/lysosomal markers. At least 30 cells per condition were analyzed across three independent experiments.

### Preparation and extraction of mouse brain MV

For brain MV isolation, brain tissue was homogenized in ice-cold modified PBS as described [72]. Brain homogenates with an equal volume of 40% Ficoll (45002020, Fisher Scientific LLC) were centrifuged to get the MV-enriched pellets. MVs were further washed and processed for WB and IP as described [73]. For WB the MVs were extracted in 1% (w/v) SDS buffer (50 mM Tris-HCl (pH 7.4), 150 mM NaCl, and 1% (w/v) SDS with protease and phosphatase inhibitors (42). For IP MVs were extracted in 1% (v/v) Triton X-100 buffer (10 mM HEPES, 2 mM CaCl_2_, 150 mM NaCl, 0.02% NaN_3_; pH 7.4) containing protease and phosphatase inhibitors [34].

### Culture of pCEC

pCECs from WT, I213T and M146V heterozygous and homozygous KI mouse brains at embryonic day 15.5 (E15.5) were isolated and purity was assessed as previously described [33].

### Microcarrier bead sprouting assay

In vitro sprouting and quantitation were performed as described [33]. WT or PS1 FAD pCECs were stimulated with vehicle (PBS) or 10ng/ml VEGF-A in vehicle [74] in the presence of vehicle (DMSO) or 200nm RO4929097 in vehicle as indicated in the figures. Number of capillary sprouts was measured as in Ref 33 in 15 microcarrier beads. At least three biological replicates (cell cultures from different animals) were analyzed. Refer to the figure legend for *n* corresponding to number of biological replicates.

### Tube formation assay

Tube formation assay was performed as previously described, with minor modifications [33]. Growth factor-reduced Matrigel (Corning) was added to µ-Slide 15-well 3D chambers (ibidi) and allowed to solidify at 37°C. Endothelial cells (1 × 10⁴ cells per well) were seeded onto the Matrigel in endothelial growth medium containing 1% (v/v) FBS with either vehicle (PBS) or 10 ng/ml VEGF-A [74] and bEnd.3 cells were stimulated with 50ng/ml VEGF-A [75] in the presence of vehicle (DMSO) or 200nm RO4929097 as indicated in the figure legends. Expression of PS1 FAD mutants or VCTF1-Flag is indicated in the figure legends. For quantification, at least nine images were acquired per condition for each independent experiment as in [33].

### Scratch Wound Healing Assay for Cell Migration

WT or PS1 FAD pCECs were cultured until a confluent monolayer was established. A linear scratch was created across the monolayer using a sterile 10-µL pipette tip as described [76]. Cells were gently washed to remove debris and then incubated in endothelial growth medium containing 1% (v/v) FBS, with vehicle (PBS) or VEGF-A (10 ng/ml) in the presence or absence of vehicle (DMSO) or 200nm RO4929097 as indicated in the figures, at 37 °C for 8–10 h. Phase-contrast images were captured immediately after scratching (0 h) and again after 8–10 h of stimulation. Wound closure was quantified using the Wound Healing Tool in ImageJ by measuring the wound area before and after VEGF-A stimulation. Cell migration was expressed as the percentage of wound closure, calculated as the percentage of wound closure (=initial wound area − remaining wound area) ÷ initial wound area × 100.

### Protein isolation, IP, and WB

For IP cell cultures were rinsed twice with PBS and lysed in 1% (v/v) Triton X-100 buffer as described [33]. IPs were carried out using specific antibodies and antigens, complexes were precipitated with Dynabeads protein A or G (Invitrogen) and detected on WB as described [33]. For WB cells were extracted in 1% (w/v) SDS buffer and proteins were detected on WB as described [33]. Semi-quantification of protein detection was done by image analysis with ImageJ.

### Immunocytochemistry (ICC)

pCECs were isolated from WT or WT/M146V or I213T mice. 50,000 cells were seeded on circular micro-coverglasses (18 mm diameter, VWR, #48380-046) coated with rat tail collagen type I (Corning, #354236). After 24 hours, cells were starved for 20 hours in a medium containing 1% (v/v) FBS. Some WT cells were pre-treated with 200nm RO4929097 overnight. Cells were then treated with 20ng/ml VEGF-A for the indicated times. Following treatment, cells were fixed with 4% (w/v) paraformaldehyde in PBS for 10 minutes at RT, permeabilized with 0.3% (v/v) Triton X-100 in PBS for 5 minutes, and blocked with 5% (v/v) newborn goat serum (Gibco, #16210064) in PBS for 1 hour at RT. The cells were then incubated overnight at 4°C with primary antibodies: anti-VEGFR2 (D5B1) at dilution 1:100 and anti-Rab5 or anti-Rab7 or anti-LAMP2 at dilution 1:300 in blocking solution. Following three times wash with PBS cells were incubated with secondary antibodies: Alexa Fluor 488-conjugated goat anti-rabbit IgG and Alexa Fluor 568-conjugated goat anti-mouse IgG at dilution 1:300 for 1 hour at RT. Nuclei were stained with 10 μg/ml Hoechst 33342 (ThermoFisher Scientific, H3570) for 5 minutes. Coverslips were mounted on slides using ProLong Diamond Antifade Mountant (Invitrogen, P36965). Images were acquired and analyzed as described above.

### In vitro γ-secretase assay

Membranes were prepared for In vitro γ-secretase assay as described [40,41] and cleavage of VEGFR2 was stimulated by VEGF-A treatment (20ng/ml) for 30 minutes in the presence and absence of 200nm RO4929097. C-terminal fragments (CTF1 and CTF2) of VEGFR2 were detected with WB as described [40,41].

### Transient transfection and transduction of cells

Transfection: HEK293T cells (ATCC) were transiently transfected with plasmids using Lipofectamine 3000 (Invitrogen), following the manufacturer’s protocol. The following constructs were used FCbAIGW empty vector (Addgene), wild-type PS1 (WT-PS1), and PS1 mutant constructs M146V, I213T, E280A, A246E, E120K, L166P, and G384A, subcloned into the FCbAIGW vector (GenScript). VEGFR2-Myc in pCMV3 vector (Sinobiological), VEGFR2-CTF1-Flag (VCTF1-Flag) and EphB2-CTF1-Flag cloned in the FCbAIGW vector (GenScript). Transduction: Transductiion using lentivirus was performed to overexpress WT PS1 and PS1 FAD mutants in HEK293T cells or VCTF1-Flag in bEnd.3 cells. All constructs in FCbAIGW vector were packaged using second-generation lentiviral plasmids psPAX2 and pMD2.G with Lipofectamine 3000 as described [77]. Viral supernatants were collected 48 hours after transfection and concentrated using the Lenti-X Concentrator (Takara Bio) according to the manufacturer’s instructions. Following transduction, GFP-positive cells were isolated by fluorescence-activated cell sorting (FACS) and expanded for subsequent experiments.

### siRNA experiments

cells were transfected with either non-targeting (NT) siRNA or ON-TARGETplus Mouse PS1 siRNA SMARTpool (L-004998-00-0005; Horizon Discovery Biosciences) using DharmaFECT 1 Transfection Reagent (T-2001-02; Horizon Discovery Biosciences) according to the manufacturer’s protocol as described [34]. In some experiments, PS1-knockdown HEK293T cells were transfected the following day with VEGFR2-Myc as described above.

### Luciferase assay

A dual luciferase assay (Promega) was performed in HEK293T cells stably expressing WT or PS1 FAD mutants (see Suppl. Fig. 2) transiently co-transfected with plasmids expressing VEGFR2-GAL4/VP16 (GenScript), 5X GAL-4-TATA-luciferase (reporter, Addgene), and Renilla Luciferase (internal control, Addgene) using Lipofectamine 3000. Luciferase was measured in cell extracts using the Dual Luciferase Reporter assay Kit (Promega) and a Varioskan Lux luminometer (ThermoScientific) according to manufacturer’s protocol.

### Bimolecular Fluorescence Complementation Analysis (BiFc)

HEK293T cells were co-transfected with cDNA expressing the BiFc fusion proteins VEGFR2-Venus(1-172) and VEGFR2-Venus(155-239) (GenScript) in pcDNA3.1 vector. For VEGFR2-Venus(1-172) the linker RSIAT was used to connect VEGFR2 with Venus and for VEGFR2-Venus(155-239) the linker was RPACKIPNDLKQKVMNH [78]. 24 hours later fluorescent and phase contrast images were captured at 20X using an Olympus IX70 microscope with Jenoptik KAPELLA. The number of positive cells counted in each field was used for quantification using ImageJ.

### Tissue processing for cryosectioning

Adult mice were anesthetized and sacrificed by transcardial perfusion with cold 4% paraformaldehyde in PBS. Following perfusion, brains were carefully removed and post-fixed in 4% paraformaldehyde for 24 h at 4 °C, transferred to 30% sucrose in PBS for cryosectioning, and stored at 4 °C until sectioning. Coronal brain sections were cut using a cryostat at a thickness of 30 µm for neuronal quantification and 40 µm for vascular staining. Sections were stored in 0.02% sodium azide in PBS at 4 °C until further use.

### Immunohistochemistry (IHC)

For Col IV staining, mouse brain sections were processed as we described [33] and incubated with an anti-Col IV (AB756P) and in experiments including MCAO with anti-GFAP antibodies for 48 hours at 4 °C, followed by incubation with goat anti-Rabbit IgG (H+L) Alexa Fluor 488 and goat anti-Chicken IgY (H+L) Alexa Fluor 555 secondary antibodies respectively for 1 hour at RT. Nuclei were counterstained with Hoechst for 5 minutes at RT. Sections were mounted on glass slides for imaging. For neuronal staining in ischemic lesions, brain sections were co-labeled with NeuN and GFAP antibodies followed by incubation with goat anti-Rabbit IgG (H+L) Alexa Fluor 488 and goat anti-Chicken IgY (H+L) Alexa Fluor 555 secondary antibodies respectively for 1 hour at RT to quantify neuronal survival in ipsilateral (lesioned) and contralateral regions. The staining protocol was similar to that described above, except that pepsin digestion was omitted, and primary antibody incubation was performed overnight at 4 °C.

### Human brain tissue

For WB brain tissue extracts were prepared in 1% (w/v) SDS buffer as described above from twelve PS1 FAD patients, each carrying a different PS1 mutation and twelve age-matched non-demented controls (Table 1). For IP brain tissue from the patients mentioned above was extracted in 1% (v/v) Triton X-100 buffer as above. For IHC tissue blocks from normal control (CT) brains or brains expressing PS1 FAD mutants (Table 2) were cryoprotected through a graded sucrose series in phosphate-buffered saline (PBS), sequential incubation in 10%, 20%, and 30% (w/v) sucrose at 4°C until each tissue block sank completely. 40μm-thick sections were processed through antigen retrieval using citrate buffer (pH 6; at 95°C for 10 min) and Tris-EDTA (pH 9; at 95°C for 10 min). Sections were then treated with pepsin (2.5 mg/ml in 1.5% HCl) for 10 min at 37°C to further improve antigen accessibility and permeabilized with 1% (v/v) Triton X-100 for 15 min, and blocked with 0.3% (v/v) Triton X-100, 5% (v/v) normal goat serum solution. Sections were incubated with anti-Col IV antibody (ab6586) at dilution 1:200 for 48 h at 4°C followed by incubation with secondary antibody goat anti-rabbit Alexa 488 at dilution 1:500 for 2 h at RT. Human brain tissue was obtained from brain banks of Mayo Clinic, University of Washington in St. Louis and NIH Neurobiobank.

### Behavioral testing

The behavioral tests include two memory and one locomotion tests, performed in the following order before and after MCAO: to asses functional cognition the Novel Object Recognition (NOR) and the Y maze tests were used. NOR test examines recognition memory. It is expressed as % discrimination index calculated as (time spent exploring the novel object – time exploring the familiar object)/(total time exploring both novel + familiar objects). Y maze test examines spatial memory and is presented as spontaneous alternations (%). Following these tests, the rotarod test was used to asses motor coordination measured as latency to fall. All tests were performed as we described [33].

### Statistical analysis

Statistical significance testing was performed with GraphPad Software. For group comparisons one- or two-way analysis of variance (ANOVA) with post hoc pairwise Tukey’s test were used as described [33]. All statistical tests are two-sided unless otherwise specified in the figure legend. *p* < 0.05 was considered significant (\**p* < 0.05, \*\**p* < 0.01, \*\*\**p* < 0.001, \*\*\*\**p* < 0.0001).

## Acknowledgments

1.We are grateful to the UW BRaIN Center and ADRC tissue donation program for samples used in this study supported by UW ADRC grant AG66509 and AG066567. Additional tissue was obtained from Florida’s AD Initiative, the Mayo Clinic ADRC supported by NIA P30AG062677, the Mayo Clinic Brain Bank (IRB25-007490) and the DIAN bank at WU supported by grant U19AG03238

2. Microscopy was performed at the Microscopy and Advanced Imaging CoRE at the Icahn School of Medicine at Mount Sinai.

3. We are grateful to Danielle Kahan and Glikeria Tzikas for excellent technical work in many experiments.

4. This work was supported by NIA grants 2R56AG008200 (NKR); 2R01AG008200 (NKR); and 2R01NS047229 (AG and NKR) and the AP Slaner family award to NKR. RP was also supported by the REC and Developmental Awards of P30AG066514 (NKR).

---

**Supplementary Figure 1.**
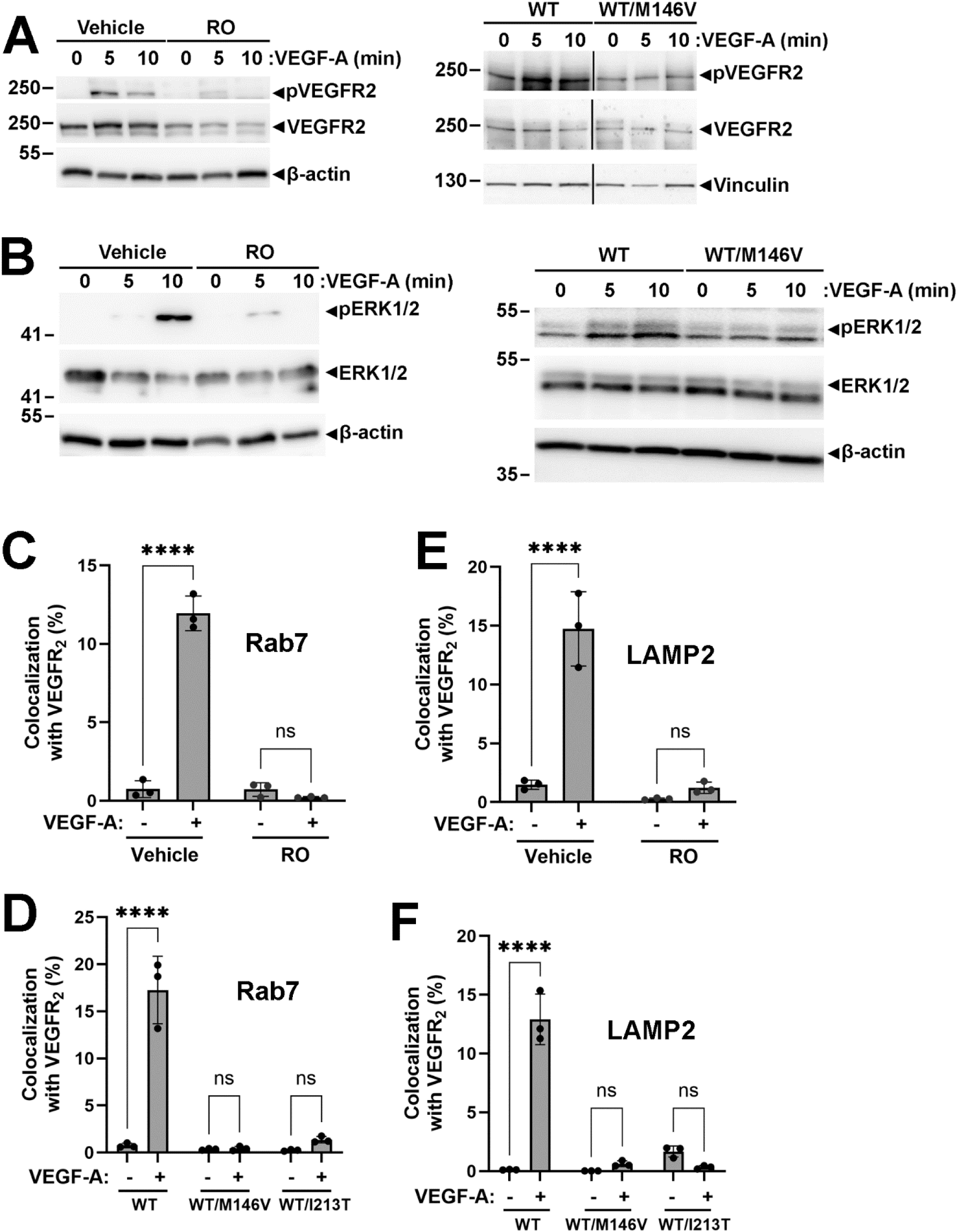
A-B: Cell culture expression of HTRZ PS1M146V or RO treatment suppress activation of pVEGFR2 and pERK1/2. Left: WT pCECs were treated with vehicle or RO and then stimulated with vehicle or VEGF-A as in 1F for indicated time periods. Cell extracts were prepared in SDS buffer as in Methods and proteins detected on WBs are shown at right of Figure. β-actin: loading control. **Right:** pCECs from WT or PS1 FAD mutant WT/M146V-expressing mice were treated with vehicle or VEGF-A as in A-B left. Cell extracts were prepared and proteins detected on WBs are shown as above. Vinculin or β-actin: loading control. Experiments performed at least twice. **C-F: PS1 FAD mutants and RO GSI decrease VEGFR2 trafficking to late endosomes and lysosomes. (C,E),** WT pCECs were treated with vehicle or RO as in 1F and then treated for 30 min with vehicle or VEGF-A as in A-B. Cell extracts were co-immunostained with anti- VEGFR2 antibodies and antibodies against late endosome marker Rab7 (C) or lysosomal marker LAMP2 (E). Co-localization of VEGFR2 with Rab7 or LAMP2 was measured and quantified as in Figs.1F-G. Graphs represent percent of VEGFR2 co-localized with Rab7 or LAMP2. **(D,F),** pCECs from WT or mutant WT/M146V- or WT/I213T-expressing mice were stimulated for 30 min with vehicle or VEGF-A as in A-B. Cells were co-stained with anti-VEGFR2 antibodies and antibodies against late endosome marker Rab7 (D) or lysosomal marker LAMP2 (F). Co-localization of VEGFR2 with Rab7 or LAMP2 was measured and quantified as in Figs. 1F-G. Graphs represent percent of VEGFR2 co-localized with Rab7 or LAMP2. **C-F:** data shown as Mean ± S.E. from three independent experiments. Statistical analysis was performed using two-way ANOVA followed by Tukey post-hoc test. ns = not significant, ****p<0.0001.

**Supplementary Figure 2.**
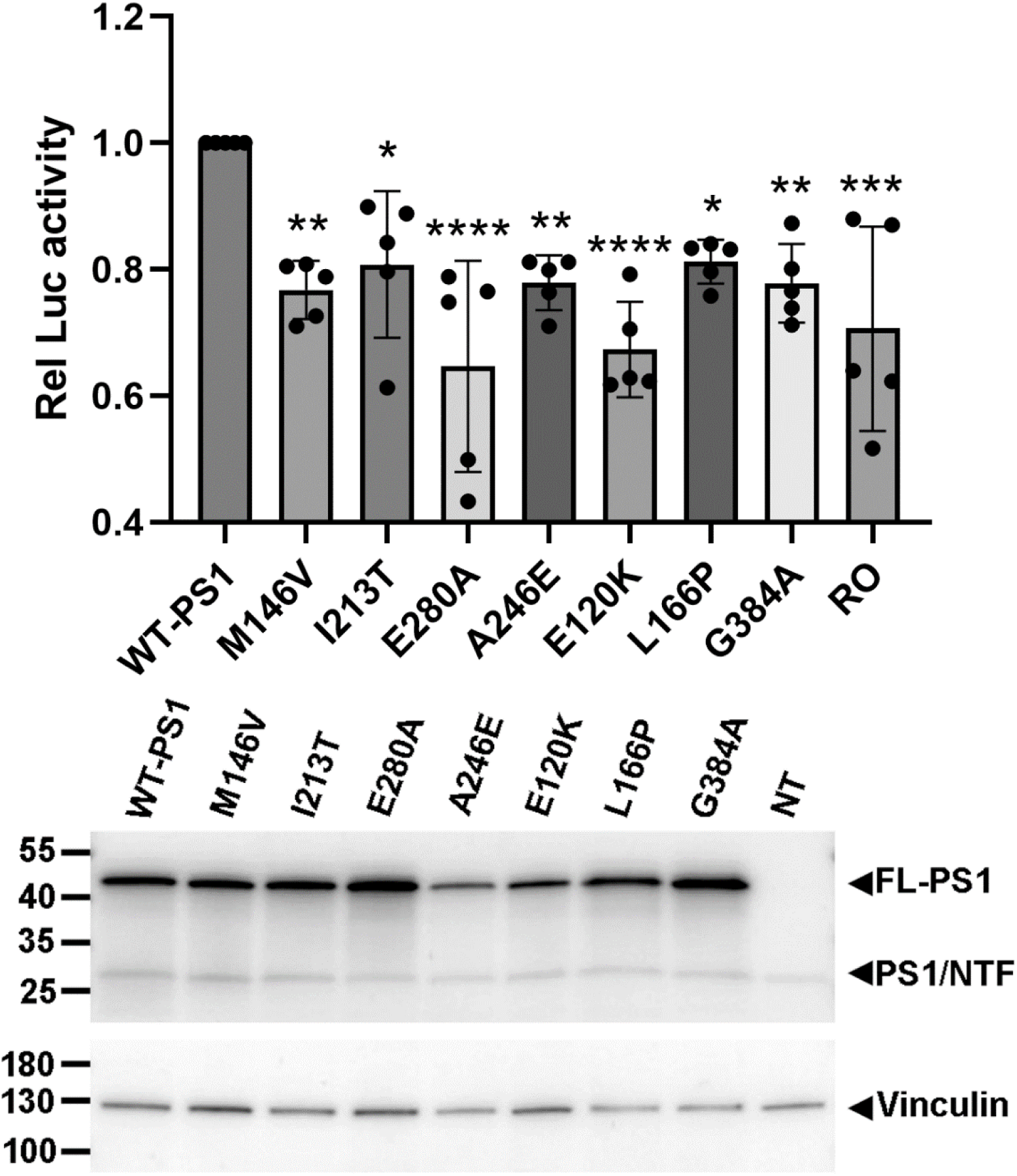
PS1 FAD mutants decrease γ-secretase processing of VEGFR2: (Upper), a dual luciferase assay (Promega) was performed as in Methods in HEK293T cells co-expressing VEGFR2-GAL4/VP16 and either WT PS1 or one of each of PS1 FAD mutants indicated in the Figure. Cells expressing WT PS1 were treated with RO as in 1F. Graph represents the γ-secretase-mediated cleavage of VEGFR2-GAL4/VP16, expressed as relative luciferase activity. Bars represent mean ± S.E. Data are from five independent experiments. Statistical significance was determined by two-way ANOVA followed by Tukey post-hoc test. ns = not significant, *p<0.05, **p<0.01, ***p<0.001. **(Lower),** Representative WB shows the expression of full length PS1 (FL-PS1) and PS1/NTF in HEK293T cells transfected with either WT PS1 (WT-PS1) or one of indicated PS1 FAD mutants. For detection of PS1 and its fragments R222 antibody was used as in 3D Right. NT: not transfected. Vinculin: loading control.

**Supplementary Figure 3.**
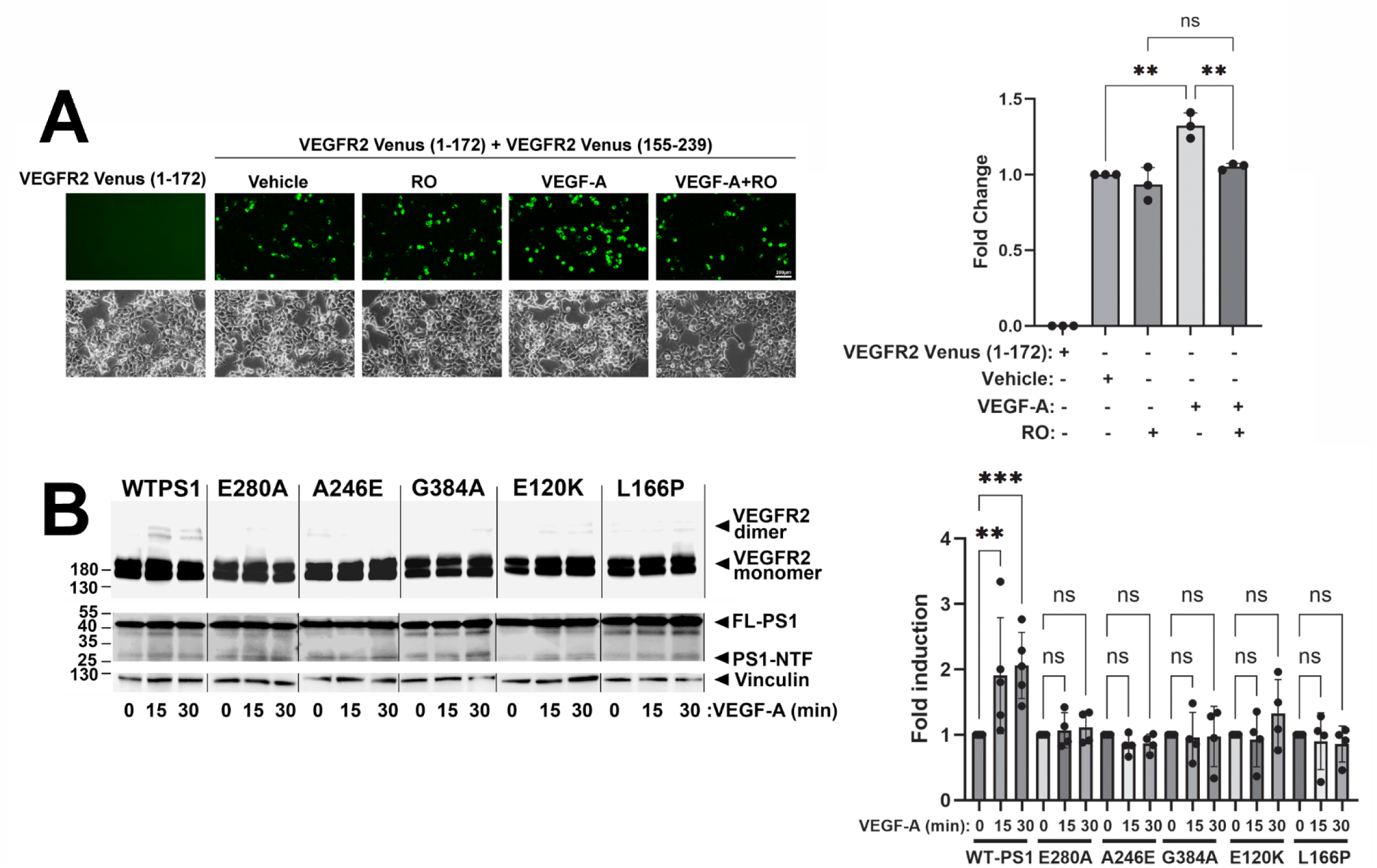
**(A),** HEK293T cells were co-transfected with both constructs expressing complementary VEGFR2-Venus fusion proteins as in Methods. Cells were treated with vehicle or VEGF-A for 30 minutes in the presence or absence of RO as in 1F and imaged as in Methods. **Left upper panel:** Green fluorescence indicates VEGFR2 homodimer formation. Scale bar: 200μm. **Left lower panel:** Corresponding phase contrast images of cell cultures. **Right:** Graph shows fold induction of VEGFR2 homodimer formation based on fluorescent intensity measured using ImageJ as in Methods. **(B),** HEK293T cells were co-transfected with VEGFR2-Myc in pCMV3 vector and either WT PS1 or one of the PS1 FAD mutants indicated in Figure in FCbAIGW. Cells were treated with vehicle or VEGF-A for 0, 15 or 30 mins and then extracted in SDS buffer as in Methods. **Left:** VEGFR2 dimers and monomers were detected on WB with anti-VEGFR2 antibody (OTI12C1). PS1 (FL-PS1) and PS1/NTF were detected with R222. Vinculin: loading control. **Right:** Graph shows fold change in the VEGFR2 dimer to monomer ratio. For statistical analysis, two-way ANOVA followed by Tukey post-hoc test was used. ns = not significant, **p<0.01, ***p<0.001.

**Supplementary Figure 4.**
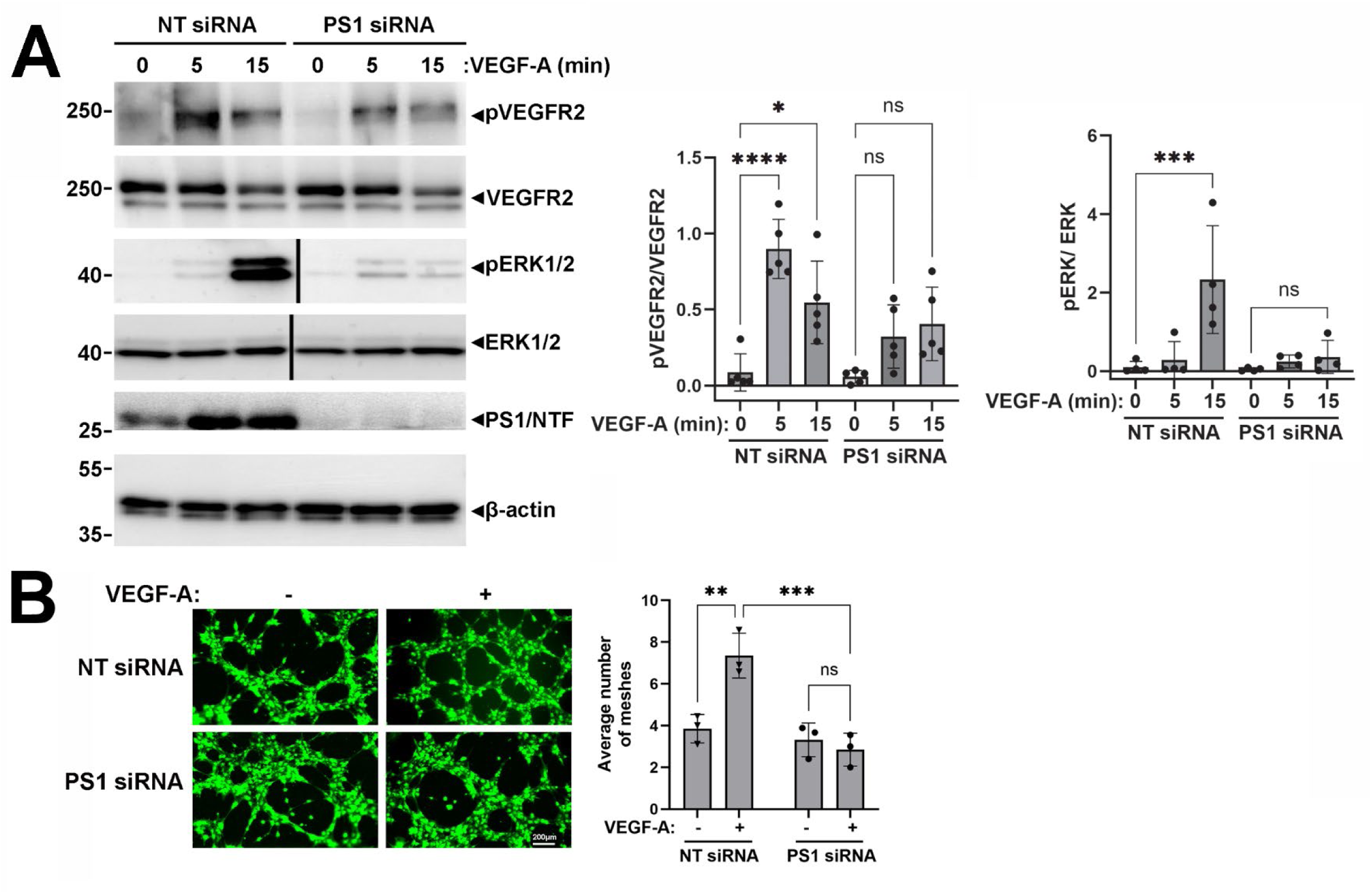
**(A),** bEnd3 cells transfected with either anti-PS1 siRNA or control non-targeting siRNA (NT siRNA) as in Methods were treated with VEGF-A (50ng/ml in PBS) for the indicated times and extracted in SDS buffer. **Left:** p-VEGFR2 (Tyr 1175), VEGFR2, p-ERK1/2 and ERK1/2 were detected on WB using specific antibodies (Cell Signaling Technology) and PS1/NTF using R222 antibody. β-actin: loading control. **Right:** Graphs show the relative levels of p-VEGFR2 (Tyr 1175) or p-ERK1/2 normalized to total VEGFR2 or ERK1/2 respectively. **(B),** bEnd3 cells transfected with either anti-PS1 siRNA or NT siRNA were plated in Matrigel and treated with vehicle or VEGF-A as in Suppl. 4A. **Left:** Representative photomicrographs show Calcein-AM-labelled tube-like structures (green). **Right:** Graph shows the quantification of tube formation represented as average number of loops/meshes per field. For statistical analysis, two-way ANOVA followed by Tukey post-hoc test was used. ns = not significant, *p<0.05, **p<0.01, ***p<0.001, ****p<0.0001.

**Supplementary Figure 5.**
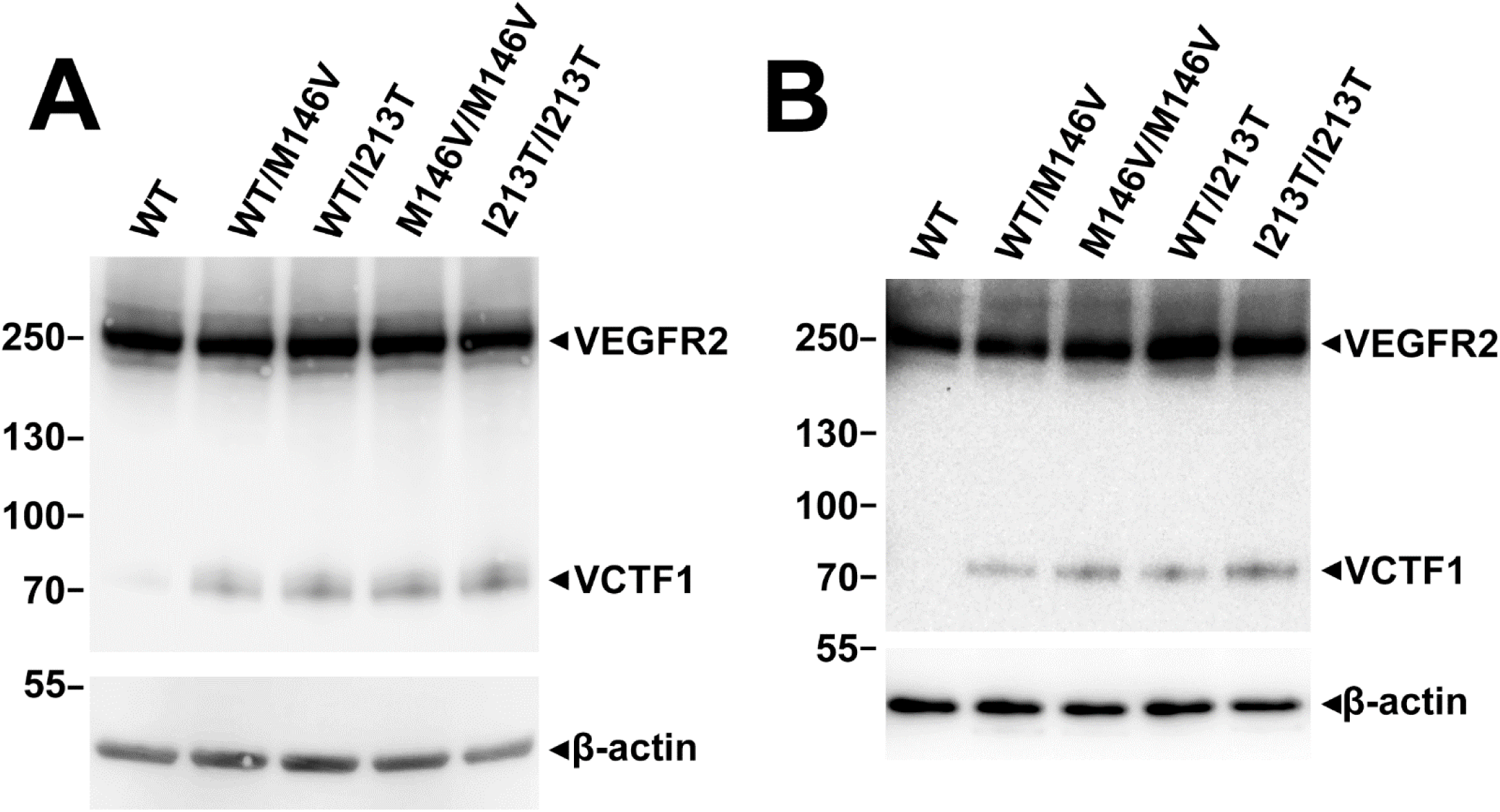
**(A),** pCECs from WT or HTRZ (WT/M146V or WT/I213T) or HMZG (M146V/M146V or I213T/I213T) mice were extracted in SDS buffer as in Methods. VEGFR2 and VCTF1 were detected on WBs of cell extracts as in 3B with the anti-VEGFR2 antibody (ab39256). β-actin: loading control. **(B),** Brain MVs were isolated from adult WT mice or mice HTRZ or HMZG for PS1 FAD mutants M146V or I213T (see Suppl. 5A). VEGFR2 and VCTF1 were detected in MV extracts on WB as in Suppl. 5A with anti-VEGFR2 antibody (ab39256). β-actin: loading control.

**Supplementary Figure 6.**
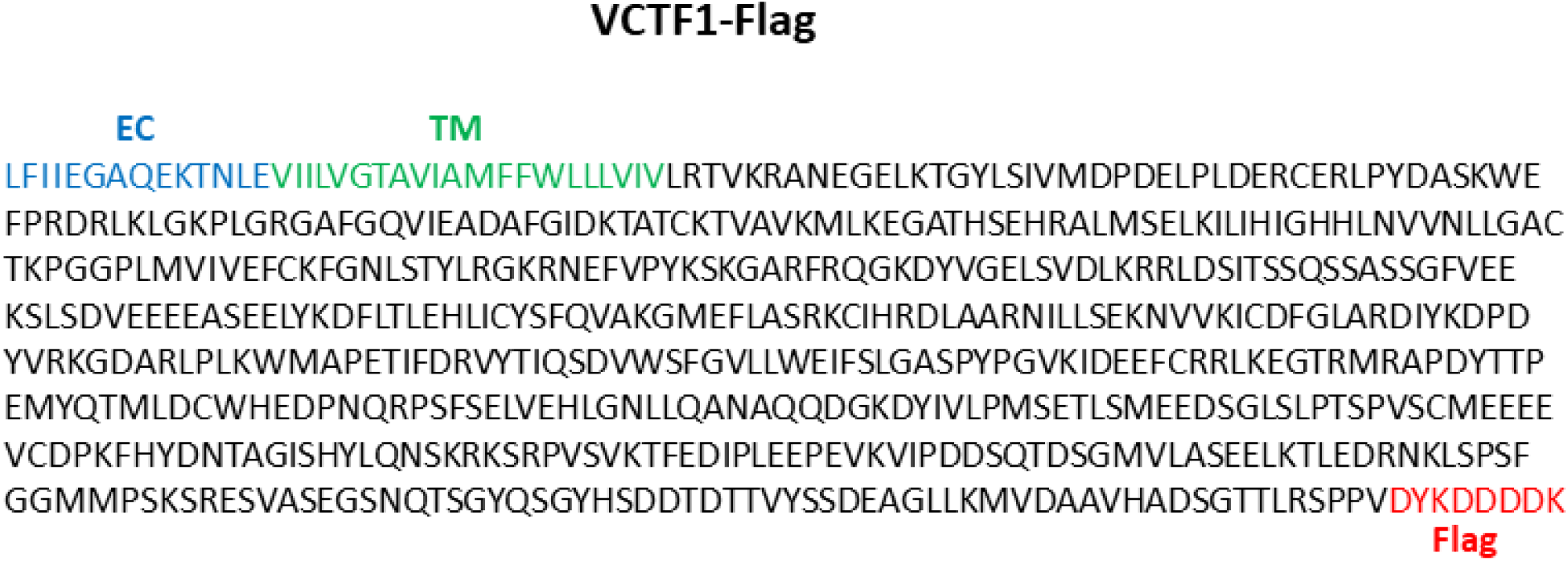
Sequence of VCTF1-Flag: VCTF1-Flag peptide includes a short extracellular segment (EC, blue), the transmembrane region (TM, green) and the cytoplasmic region of mouse VEGFR2 (GenBank: NP_001390070) followed by a Flag tag (Flag, red).

## References

1. Kapasi, A., DeCarli, C., and Schneider, J. A. (2017) Impact of multiple pathologies on the threshold for clinically overt dementia. Acta Neuropathol 134, 171–186

2. James, B. D., Wilson, R. S., Boyle, P. A., Trojanowski, J. Q., Bennett, D. A., and Schneider, J. A. (2016) TDP-43 stage, mixed pathologies, and clinical Alzheimer’s-type dementia. Brain : a journal of neurology 139, 2983–2993

3. Jellinger, K. A., and Attems, J. (2015) Challenges of multimorbidity of the aging brain: a critical update. J Neural Transm (Vienna) 122, 505–521

4. Toledo, J. B., Arnold, S. E., Raible, K., Brettschneider, J., Xie, S. X., Grossman, M., et al. (2013) Contribution of cerebrovascular disease in autopsy confirmed neurodegenerative disease cases in the National Alzheimer’s Coordinating Centre. Brain : a journal of neurology 136, 2697–2706

5. Eisenmenger, L. B., Peret, A., Famakin, B. M., Spahic, A., Roberts, G. S., Bockholt, J. H., et al. (2023) Vascular contributions to Alzheimer’s disease. Transl Res 254, 41–53

6. Aguero-Torres, H., Kivipelto, M., and von Strauss, E. (2006) Rethinking the dementia diagnoses in a population-based study: what is Alzheimer’s disease and what is vascular dementia?. A study from the kungsholmen project. Dementia and geriatric cognitive disorders 22, 244–249

7. Kovari, E., Gold, G., Herrmann, F. R., Canuto, A., Hof, P. R., Bouras, C., et al. (2007) Cortical microinfarcts and demyelination affect cognition in cases at high risk for dementia. Neurology 68, 927–931

8. Buee, L., Hof, P. R., Bouras, C., Delacourte, A., Perl, D. P., Morrison, J. H., et al. (1994) Pathological alterations of the cerebral microvasculature in Alzheimer’s disease and related dementing disorders. Acta Neuropathol 87, 469–480

9. Lee, H., Kim, K., Lee, Y. C., Kim, S., Won, H. H., Yu, T. Y., et al. (2020) Associations between vascular risk factors and subsequent Alzheimer’s disease in older adults. Alzheimers Res Ther 12, 117

10. Mohanty, R., Wheatley, S., Chiotis, K., Marseglia, A., Westman, E., and Alzheimer’s Disease Neuroimaging Initiative, C. (2025) Distinct cerebrovascular pathways underlying Alzheimer’s disease-related neurodegeneration. Acta Neuropathol 150, 64

11. Kalaria, R. N., and Sepulveda-Falla, D. (2021) Cerebral Small Vessel Disease in Sporadic and Familial Alzheimer Disease. The American journal of pathology 191, 1888–1905

12. Szpak, G. M., Lewandowska, E., Wierzba-Bobrowicz, T., Bertrand, E., Pasennik, E., Mendel, T., et al. (2007) Small cerebral vessel disease in familial amyloid and non-amyloid angiopathies: FAD-PS-1 (P117L) mutation and CADASIL. Immunohistochemical and ultrastructural studies. *Folia neuropathologica / Association of Polish Neuropathologists and Medical Research Centre*, Polish Academy of Sciences 45, 192–204

13. Littau, J. L., Velilla, L., Hase, Y., Villalba-Moreno, N. D., Hagel, C., Drexler, D., et al. (2022) Evidence of beta amyloid independent small vessel disease in familial Alzheimer’s disease. Brain Pathol 32, e13097

14. Bailey, T. L., Rivara, C. B., Rocher, A. B., and Hof, P. R. (2004) The nature and effects of cortical microvascular pathology in aging and Alzheimer’s disease. Neurological research 26, 573–578

15. Iturria-Medina, Y., Sotero, R. C., Toussaint, P. J., Mateos-Perez, J. M., Evans, A. C., and Alzheimer’s Disease Neuroimaging, I. (2016) Early role of vascular dysregulation on late-onset Alzheimer’s disease based on multifactorial data-driven analysis. Nat Commun 7, 11934

16. Nielsen, R. B., Egefjord, L., Angleys, H., Mouridsen, K., Gejl, M., Moller, A., et al. (2017) Capillary dysfunction is associated with symptom severity and neurodegeneration in Alzheimer’s disease. Alzheimers Dement 13, 1143–1153

17. Govindpani, K., Vinnakota, C., Waldvogel, H. J., Faull, R. L., and Kwakowsky, A. (2020) Vascular dysfunction in Alzheimer’s disease: a biomarker of disease progression and a potential therapeutic target. Neural Regen Res 15, 1030–1032

18. Grammas, P. (2011) Neurovascular dysfunction, inflammation and endothelial activation: implications for the pathogenesis of Alzheimer’s disease. J Neuroinflammation 8, 26

19. Hatakeyama, M., Ninomiya, I., and Kanazawa, M. (2020) Angiogenesis and neuronal remodeling after ischemic stroke. Neural Regen Res 15, 16–19

20. Kuzma, E., Lourida, I., Moore, S. F., Levine, D. A., Ukoumunne, O. C., and Llewellyn, D. J. (2018) Stroke and dementia risk: A systematic review and meta-analysis. Alzheimers Dement 14, 1416–1426

21. Pluta, R., Januszewski, S., and Czuczwar, S. J. (2021) Brain Ischemia as a Prelude to Alzheimer’s Disease. Front Aging Neurosci 13, 636653

22. Lange, C., Storkebaum, E., de Almodovar, C. R., Dewerchin, M., and Carmeliet, P. (2016) Vascular endothelial growth factor: a neurovascular target in neurological diseases. Nat Rev Neurol 12, 439–454

23. Simons, M., Gordon, E., and Claesson-Welsh, L. (2016) Mechanisms and regulation of endothelial VEGF receptor signalling. Nature reviews. Molecular cell biology 17, 611–625

24. Sarabipour, S., Ballmer-Hofer, K., and Hristova, K. (2016) VEGFR-2 conformational switch in response to ligand binding. Elife 5, e13876

25. Koh, B. I., Lee, H. J., Kwak, P. A., Yang, M. J., Kim, J. H., Kim, H. S., et al. (2020) VEGFR2 signaling drives meningeal vascular regeneration upon head injury. Nat Commun 11, 3866

26. Benedictus, M. R., Leeuwis, A. E., Binnewijzend, M. A., Kuijer, J. P., Scheltens, P., Barkhof, F., et al. (2017) Lower cerebral blood flow is associated with faster cognitive decline in Alzheimer’s disease. Eur Radiol 27, 1169–1175

27. Kalaria, R. N., Cohen, D. L., Premkumar, D. R., Nag, S., LaManna, J. C., and Lust, W. D. (1998) Vascular endothelial growth factor in Alzheimer’s disease and experimental cerebral ischemia. Brain Res Mol Brain Res 62, 101–105

28. Tang, H., Mao, X., Xie, L., Greenberg, D. A., and Jin, K. (2013) Expression level of vascular endothelial growth factor in hippocampus is associated with cognitive impairment in patients with Alzheimer’s disease. Neurobiol Aging 34, 1412–1415

29. Swendeman, S., Mendelson, K., Weskamp, G., Horiuchi, K., Deutsch, U., Scherle, et al. (2008) VEGF-A stimulates ADAM17-dependent shedding of VEGFR2 and crosstalk between VEGFR2 and ERK signaling. Circulation research 103, 916–918

30. Wolfe, M. S. (2019) Structure and Function of the gamma-Secretase Complex. Biochemistry 58, 2953–2966

31. Barthet, G., Georgakopoulos, A., and Robakis, N. K. (2012) Cellular mechanisms of gamma-secretase substrate selection, processing and toxicity. Progress in neurobiology 98, 166–175

32. Barthet, G., Shioi, J., Shao, Z., Ren, Y., Georgakopoulos, A., and Robakis, N. K. (2011) Inhibitors of gamma-secretase stabilize the complex and differentially affect processing of amyloid precursor protein and other substrates. FASEB journal : official publication of the Federation of American Societies for Experimental Biology 25, 2937–2946

33. Yoon, Y., Voloudakis, G., Doran, N., Zhang, E., Dimovasili, C., Chen, L., et al. (2021) PS1 FAD mutants decrease ephrinB2-regulated angiogenic functions, ischemia-induced brain neovascularization and neuronal survival. Mol Psychiatry 26, 1996–2012

34. Al Rahim, M., Yoon, Y., Dimovasili, C., Shao, Z., Huang, Q., Zhang, E., et al. (2020) Presenilin1 familial Alzheimer disease mutants inactivate EFNB1- and BDNF-dependent neuroprotection against excitotoxicity by affecting neuroprotective complexes of N-methyl-d-aspartate receptor. Brain Commun 2, fcaa100

35. Guo, Q., Fu, W., Sopher, B. L., Miller, M. W., Ware, C. B., Martin, G. M., and Mattson, M. P. (1999) Increased vulnerability of hippocampal neurons to excitotoxic necrosis in presenilin-1 mutant knock-in mice. Nature medicine 5, 101–106

36. Nakano, Y., Kondoh, G., Kudo, T., Imaizumi, K., Kato, M., Miyazaki, J. I., et al. (1999) Accumulation of murine amyloidbeta42 in a gene-dosage-dependent manner in PS1 ’knock-in’ mice. The European journal of neuroscience 11, 2577–2581

37. Manoonkitiwongsa, P. S., Schultz, R. L., McCreery, D. B., Whitter, E. F., and Lyden, P. D. (2004) Neuroprotection of ischemic brain by vascular endothelial growth factor is critically dependent on proper dosage and may be compromised by angiogenesis. Journal of cerebral blood flow and metabolism : official journal of the International Society of Cerebral Blood Flow and Metabolism 24, 693–702

38. Norrby, K. (1998) Microvascular density in terms of number and length of microvessel segments per unit tissue volume in mammalian angiogenesis. Microvascular research 55, 43–53

39. Sharma, S., Ehrlich, M., Zhang, M., Blobe, G. C., and Henis, Y. I. (2024) NRP1 interacts with endoglin and VEGFR2 to modulate VEGF signaling and endothelial cell sprouting. Commun Biol 7, 112

40. Litterst, C., Georgakopoulos, A., Shioi, J., Ghersi, E., Wisniewski, T., Wang, R., et al. (2007) Ligand binding and calcium influx induce distinct ectodomain/gamma-secretase-processing pathways of EphB2 receptor. The Journal of biological chemistry 282, 16155–16163

41. Marambaud, P., Wen, P. H., Dutt, A., Shioi, J., Takashima, A., Siman, R., et al. (2003) A CBP binding transcriptional repressor produced by the PS1/epsilon-cleavage of N-cadherin is inhibited by PS1 FAD mutations. Cell 114, 635–645

42. Georgakopoulos, A., Litterst, C., Ghersi, E., Baki, L., Xu, C., Serban, G., et al. (2006) Metalloproteinase/Presenilin1 processing of ephrinB regulates EphB-induced Src phosphorylation and signaling. The EMBO journal 25, 1242–1252

43. Watanabe, H., and Shen, J. (2017) Dominant negative mechanism of Presenilin-1 mutations in FAD. Proceedings of the National Academy of Sciences of the United States of America 114, 12635–12637

44. Kalen, M., Heikura, T., Karvinen, H., Nitzsche, A., Weber, H., Esser, N., et al. (2011) Gamma-secretase inhibitor treatment promotes VEGF-A-driven blood vessel growth and vascular leakage but disrupts neovascular perfusion. PloS one 6, e18709

45. Cao, L., Arany, P. R., Kim, J., Rivera-Feliciano, J., Wang, Y. S., He, Z., et al. (2010) Modulating Notch signaling to enhance neovascularization and reperfusion in diabetic mice. Biomaterials 31, 9048–9056

46. Luistro, L., He, W., Smith, M., Packman, K., Vilenchik, M., Carvajal, D., et al. (2009) Preclinical profile of a potent gamma-secretase inhibitor targeting notch signaling with in vivo efficacy and pharmacodynamic properties. Cancer Res 69, 7672–7680

47. Zhang, C., Qin, S., Xie, H., Qiu, Q., Wang, H., Zhang, J., et al. (2022) RO4929097, a Selective gamma-Secretase Inhibitor, Inhibits Subretinal Fibrosis Via Suppressing Notch and ERK1/2 Signaling in Laser-Induced Mouse Model. Investigative ophthalmology & visual science 63, 14

48. Gounder, M. M., Rosenbaum, E., Wu, N., Dickson, M. A., Sheikh, T. N., D’Angelo, S. P., et al. (2022) A Phase Ib/II Randomized Study of RO4929097, a Gamma-Secretase or Notch Inhibitor with or without Vismodegib, a Hedgehog Inhibitor, in Advanced Sarcoma. Clin Cancer Res 28, 1586–1594

49. Wang, X., Bove, A. M., Simone, G., and Ma, B. (2020) Molecular Bases of VEGFR-2-Mediated Physiological Function and Pathological Role. Front Cell Dev Biol 8, 599281

50. Basagiannis, D., Zografou, S., Murphy, C., Fotsis, T., Morbidelli, L., Ziche, M., et al. (2016) VEGF induces signalling and angiogenesis by directing VEGFR2 internalisation through macropinocytosis. J Cell Sci 129, 4091–4104

51. Goodwin, A. M. (2007) In vitro assays of angiogenesis for assessment of angiogenic and anti-angiogenic agents. Microvascular research 74, 172–183

52. Smith, G. A., Fearnley, G. W., Abdul-Zani, I., Wheatcroft, S. B., Tomlinson, D. C., Harrison, M. A., et al. (2016) VEGFR2 Trafficking, Signaling and Proteolysis is Regulated by the Ubiquitin Isopeptidase USP8. Traffic 17, 53–65

53. Bertuccio, C., Veron, D., Aggarwal, P. K., Holzman, L., and Tufro, A. (2011) Vascular endothelial growth factor receptor 2 direct interaction with nephrin links VEGF-A signals to actin in kidney podocytes. The Journal of biological chemistry 286, 39933–39944

54. Gooz, M. (2010) ADAM-17: the enzyme that does it all. Crit Rev Biochem Mol Biol 45, 146–169

55. Zhou, J., Li, S., Leung, K. K., O’Donovan, B., Zou, J. Y., DeRisi, J. L., et al. (2020) Deep profiling of protease substrate specificity enabled by dual random and scanned human proteome substrate phage libraries. Proceedings of the National Academy of Sciences of the United States of America 117, 25464–25475

56. Escamilla-Ayala, A., Wouters, R., Sannerud, R., and Annaert, W. (2020) Contribution of the Presenilins in the cell biology, structure and function of gamma-secretase. Semin Cell Dev Biol 105, 12–26

57. Wiley, J. C., Hudson, M., Kanning, K. C., Schecterson, L. C., and Bothwell, M. (2005) Familial Alzheimer’s disease mutations inhibit gamma-secretase-mediated liberation of beta-amyloid precursor protein carboxy-terminal fragment. J Neurochem 94, 1189–1201

58. Peach, C. J., Mignone, V. W., Arruda, M. A., Alcobia, D. C., Hill, S. J., Kilpatrick, L. E., et al. (2018) Molecular Pharmacology of VEGF-A Isoforms: Binding and Signalling at VEGFR2. Int J Mol Sci 19

59. Ahmadova, Z., Yagublu, V., Forg, T., Hajiyeva, Y., Jesenofsky, R., Hafner, M., et al. (2014) Fluorescent resonance energy transfer imaging of VEGFR dimerization. Anticancer Res 34, 2123–2133

60. Zhou, R., Yang, G., and Shi, Y. (2017) Dominant negative effect of the loss-of-function gamma-secretase mutants on the wild-type enzyme through heterooligomerization. Proceedings of the National Academy of Sciences of the United States of America 114, 12731–12736

61. Robakis, N. K., and Georgakopoulos, A. (2014) Allelic interference: a mechanism for trans-dominant transmission of loss of function in the neurodegeneration of familial Alzheimer’s disease. Neurodegener Dis 13, 126–130

62. Tosto, G., Bird, T. D., Bennett, D. A., Boeve, B. F., Brickman, A. M., Cruchaga, C., et al. (2016) The Role of Cardiovascular Risk Factors and Stroke in Familial Alzheimer Disease. JAMA Neurol 73, 1231–1237

63. Sun, Y., Jin, K., Xie, L., Childs, J., Mao, X. O., Logvinova, A., et al. (2003) VEGF-induced neuroprotection, neurogenesis, and angiogenesis after focal cerebral ischemia. J Clin Invest 111, 1843–1851

64. Yang, J., Yao, Y., Chen, T., and Zhang, T. (2014) VEGF ameliorates cognitive impairment in in vivo and in vitro ischemia via improving neuronal viability and function. Neuromolecular Med 16, 376–388

65. Kilic, U., Kilic, E., Jarve, A., Guo, Z., Spudich, A., Bieber, K., et al. (2006) Human vascular endothelial growth factor protects axotomized retinal ganglion cells in vivo by activating ERK-1/2 and Akt pathways. J Neurosci 26, 12439–12446

66. Franberg, J., Karlstrom, H., Winblad, B., Tjernberg, L. O., and Frykman, S. (2010) gamma-Secretase dependent production of intracellular domains is reduced in adult compared to embryonic rat brain membranes. PloS one 5, e9772

67. Shen, J., Bronson, R. T., Chen, D. F., Xia, W., Selkoe, D. J., and Tonegawa, S. (1997) Skeletal and CNS defects in Presenilin-1-deficient mice. Cell 89, 629–639

68. Longa, E. Z., Weinstein, P. R., Carlson, S., and Cummins, R. (1989) Reversible middle cerebral artery occlusion without craniectomy in rats. Stroke; a journal of cerebral circulation 20, 84–91

69. Leclerc, M., Bourassa, P., Tremblay, C., Caron, V., Sugere, C., Emond, V., et al. (2023) Cerebrovascular insulin receptors are defective in Alzheimer’s disease. Brain : a journal of neurology 146, 75–90

70. Best, J. D., Jay, M. T., Otu, F., Churcher, I., Reilly, M., Morentin-Gutierrez, P., et al. (2006) In vivo characterization of Abeta(40) changes in brain and cerebrospinal fluid using the novel gamma-secretase inhibitor N-[cis-4-[(4-chlorophenyl)sulfonyl]-4-(2,5-difluorophenyl)cyclohexyl]-1,1,1-trifluoromethanesulfonamide (MRK-560) in the rat. J Pharmacol Exp Ther 317, 786–790

71. Li, W. L., Fraser, J. L., Yu, S. P., Zhu, J., Jiang, Y. J., and Wei, L. (2011) The role of VEGF/VEGFR2 signaling in peripheral stimulation-induced cerebral neurovascular regeneration after ischemic stroke in mice. Exp Brain Res 214, 503–513

72. Lee, Y. K., Uchida, H., Smith, H., Ito, A., and Sanchez, T. (2019) The isolation and molecular characterization of cerebral microvessels. Nat Protoc 14, 3059–3081

73. Paraiso, H. C., Wang, X., Kuo, P. C., Furnas, D., Scofield, B. A., Chang, F. L., et al. (2020) Isolation of Mouse Cerebral Microvasculature for Molecular and Single-Cell Analysis. Front Cell Neurosci 14, 84

74. Carpentier, G., Berndt, S., Ferratge, S., Rasband, W., Cuendet, M., Uzan, G., et al. (2020) Angiogenesis Analyzer for ImageJ - A comparative morphometric analysis of “Endothelial Tube Formation Assay” and “Fibrin Bead Assay”. Sci Rep 10, 11568

75. Jaiprasart, P., Dogra, S., Neelakantan, D., Devapatla, B., and Woo, S. (2020) Identification of signature genes associated with therapeutic resistance to anti-VEGF therapy. Oncotarget 11, 99–114

76. Liang, C. C., Park, A. Y., and Guan, J. L. (2007) In vitro scratch assay: a convenient and inexpensive method for analysis of cell migration in vitro. Nat Protoc 2, 329–333

77. Elegheert, J., Behiels, E., Bishop, B., Scott, S., Woolley, R. E., Griffiths, S. C., et al. (2018) Lentiviral transduction of mammalian cells for fast, scalable and high-level production of soluble and membrane proteins. Nat Protoc 13, 2991–3017

78. Kerppola, T. K. (2006) Design and implementation of bimolecular fluorescence complementation (BiFC) assays for the visualization of protein interactions in living cells. Nat Protoc 1, 1278–1286

